# Predicting VHH-Fc Developability from Large-Scale IgG Data

**DOI:** 10.64898/2026.08.18.745296

**Authors:** Joshua Moller, Seth Ritter, Laura Rand, Alexander Smith, Yolaine Pierre, Tim Bloomingdale, Blake Harris, Sai Karthick, Lucia Grippo, Aanal Bhatt, Jay Patel, Xiang Ao, Rebecca Bhatt, Rich Cohen, David W. Borhani, Peter M. Tessier, Ammar Arsiwala

## Abstract

The VHH-Fc antibody scaffold is an emerging therapeutic modality. No public large-scale, standardized developability VHH-Fc dataset exists. Filling that gap, we introduce GDPa5, a 160-member VHH-Fc library profiled across 10 biophysical assays on the PROPHET-Ab platform. Cross format models trained on the developability properties of 559 IgGs outperformed intra-format models trained on GDPa5 alone, which is an advantage driven by the larger scale of standardized IgG data rather than by format. The most accurately predicted properties were heparin binding (HAC, Spearman ρ=0.82), hydrophobicity (HIC, ρ=0.63), and self-association (AC-SINS, ρ=0.62), all of which are largely governed by antibody surface properties. Tabular neural networks (TabICLv2, TabPFN v2.5), applied here for the first time to antibody developability prediction, outperformed conventional modeling approaches. Adding experimental HIC and HAC measurements as model inputs improved prediction of the more complex polyreactivity liability (PR-CHO, Δρ = +0.10), supporting a tiered assay strategy that extends predictive performance while limiting experimental burden. We demonstrate through this work that IgG-trained models are a practical, data-efficient starting point for VHH-Fc developability prediction.

**Significance:** The therapeutic antibody landscape is rapidly diversifying beyond conventional IgG scaffolds, yet developability prediction tools have not kept pace. Format-specific training data are scarce for emerging scaffolds; whether models trained on IgGs can predict the developability properties of other scaffolds has not been systematically evaluated. In this work, we present GDPa5, a 160-member VHH-Fc developability dataset, and demonstrate that models trained exclusively on IgG data can predict VHH-Fc developability without VHH-Fc-specific training. The GDPa5 dataset and the predictive power of tabular neural networks establish a data-efficient modeling framework broadly applicable to emerging antibody formats.

## Introduction

Single-domain antibodies (VHHs or nanobodies) bind targets through a single immunoglobulin domain that combines high affinity with compact size (∼15 kDa), high thermostability, and the ability to access epitopes sterically occluded from conventional antibodies [Mitchell & Colwell, 2018]. Fusion of a VHH to an IgG Fc region (VHH-Fc) extends serum half-life through FcRn recycling and adds effector function, without sacrificing these favorable properties [De Vlieger et al., 2019]. The growing clinical validation of this format has accelerated its adoption. In particular, caplacizumab was the first approved VHH-based therapeutic, and multiple VHH-Fcs have since entered or are progressing through development [Duggan, 2018].

To enable economical manufacturing, stable formulation, and safe clinical use, antibodies must also possess suitable developability attributes, such as thermostability, low self-association and polyreactivity, and acceptable hydrophobicity [Jain et al., 2017]. These properties are inherent to an antibody’s amino acid sequence and structure and ideally assessed early in discovery to avoid costly late-stage attrition [Jain et al., 2023]. High-throughput experimental platforms have emerged to address this need at scale. Recent work from Ginkgo Datapoints has focused on the release of standardized, public datasets for 246 clinical-stage IgGs (GDPa1), 80 sequence diverse IgGs from paired OAS (GDPa3), and 160 bispecific antibodies (GDPa4) [Arsiwala et al., 2025; van Niekerk et al., 2026; Ritter et al., 2026]. Despite the increase in publicly available data, generalizable predictive models remain constrained by a persistent data bottleneck. The 2025 Ginkgo Datapoints Antibody Developability Competition demonstrated that even state-of-the-art algorithms overfit to training distributions and show limited out-of-distribution generalization [van Niekerk et al., 2026]. For the VHH format specifically, the Therapeutic Nanobody Profiler (TNP) recently characterized 36 clinical nanobody sequences across a panel of biophysical assays, which is a valuable clinical benchmark but at a scale insufficient for model training [Gordon et al., 2026].

Extending beyond single formats, the field has long operated under the assumption that predicting developability for a new format, such as VHH-Fcs or bispecifics, requires generating carefully curated, format-specific experimental data. This is a resource-intensive requirement that compounds the challenge as the therapeutic landscape continues to diversify. Recent work has begun to challenge this assumption. For bispecific antibodies, systematic profiling of 160 bispecifics and their 65 parental monospecific arms demonstrated that most developability properties can be predicted from parental measurements, with hydrophobicity and surface charge transferring most reliably (Spearman ρ ≈ 0.85–0.95) and thermostability requiring direct measurement on the bispecific itself (ρ < 0.4) [Ritter et al., 2026]. For scFv-to-IgG transfer, early evidence suggests that sequence embeddings can partially bridge structural formats [Xin et al., 2025]. Collectively, these observations raise a central question: when generated under standardized conditions on a common platform, can the substantial IgG developability data already available be deployed to predict the biophysical behavior of VHH-Fcs, despite their structural divergence?

Here, we address this question directly. We present GDPa5, a diverse 160-member VHH-Fc library selected from the PLAbDAb-nano database using farthest-point sampling and profiled across 10 biophysical assays on the PROPHET-Ab platform. This dataset is released as a public resource, complementing GDPa1 and GDPa3 to support cross-format modeling across the community. Using GDPa5, we establish three findings (**Figure 1)**. First, we find at this data scale, intra-format models trained solely on VHH-Fcs alone are limited by data scarcity. Second, tabular neural networks trained on a larger corpus of 559 IgGs exploit that scale to outperform format-specific VHH-Fc models in zero-shot prediction, despite the structural divergence between formats. Third, a hierarchical assay cascade, wherein measuring simple surface properties first and using them as inputs to predict complex liabilities, extends cross-format predictive performance. Together, these results establish a data-efficient framework for VHH-Fc developability prediction that leverages, rather than replaces, existing IgG datasets.

**Figure 1:**
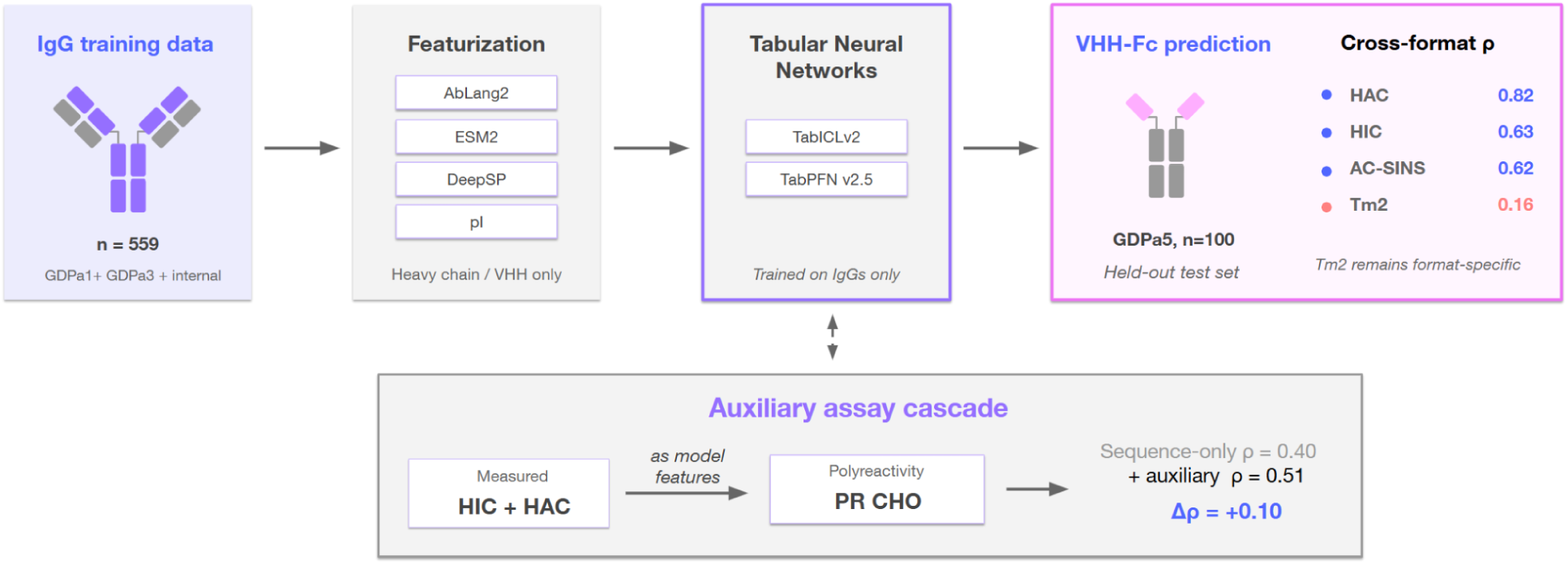
A cross-format framework for VHH-Fc developability prediction. Tabular neural networks (TabICLv2, TabPFN v2.5) were trained on a 559-antibody IgG dataset (GDPa1, GDPa3, and internal Ginkgo data) using heavy chain sequences only, featurized with protein language model embeddings (AbLang2, ESM2), spatial structural descriptors (DeepSP), and isoelectric point (pI). Trained models were applied zero-shot to GDPa5, a 160-member VHH-Fc library profiled across 10 biophysical assays on the PROPHET-Ab platform (n = 100 after filtering for ≥80% SEC monomer purity). Despite the structural divergence between formats, properties governed by localized surface features were predicted relatively well: heparin binding (HAC, ρ = 0.82), hydrophobicity (HIC, ρ = 0.63), and self-association (AC-SINS, ρ = 0.62). Thermostability does not (Tm2, ρ = 0.16) and remains a format-specific measurement. This advantage derives from the scale of available IgG data rather than the IgG format itself; at matched training set size, format-specific models remain competitive. A tiered auxiliary cascade (lower panel) extends the framework: hydrophobicity and heparin binding values, measured or predicted from sequence, serve as additional inputs and improve polyreactivity prediction (PR CHO, ρ = 0.40 → 0.51).

## Results

### GDPa5: a diverse VHH-Fc library representing clinical and broader VHH sequence space

To enable rigorous evaluation of cross-format developability prediction, we designed a VHH library, “GDPa5,” to maximize sequence diversity anchored to clinically relevant VHH sequences. We started from a pool of 4,913 sequences from PLAbDAb-nano [Gordon et al., 2025], a curated database of single-domain antibody sequences, and filtered these to give 2,306 deduplicated sequences exceeding 100 amino acids in length. From this set, we applied a length filter and farthest-point sampling (FPS), seeded with 36 clinical nanobody sequences drawn from [Gordon et al., 2026], to select 160 sequences that maximize coverage of VHH sequence diversity anchored to clinically relevant sequences (**Figure 2A**).

**Figure 2:**
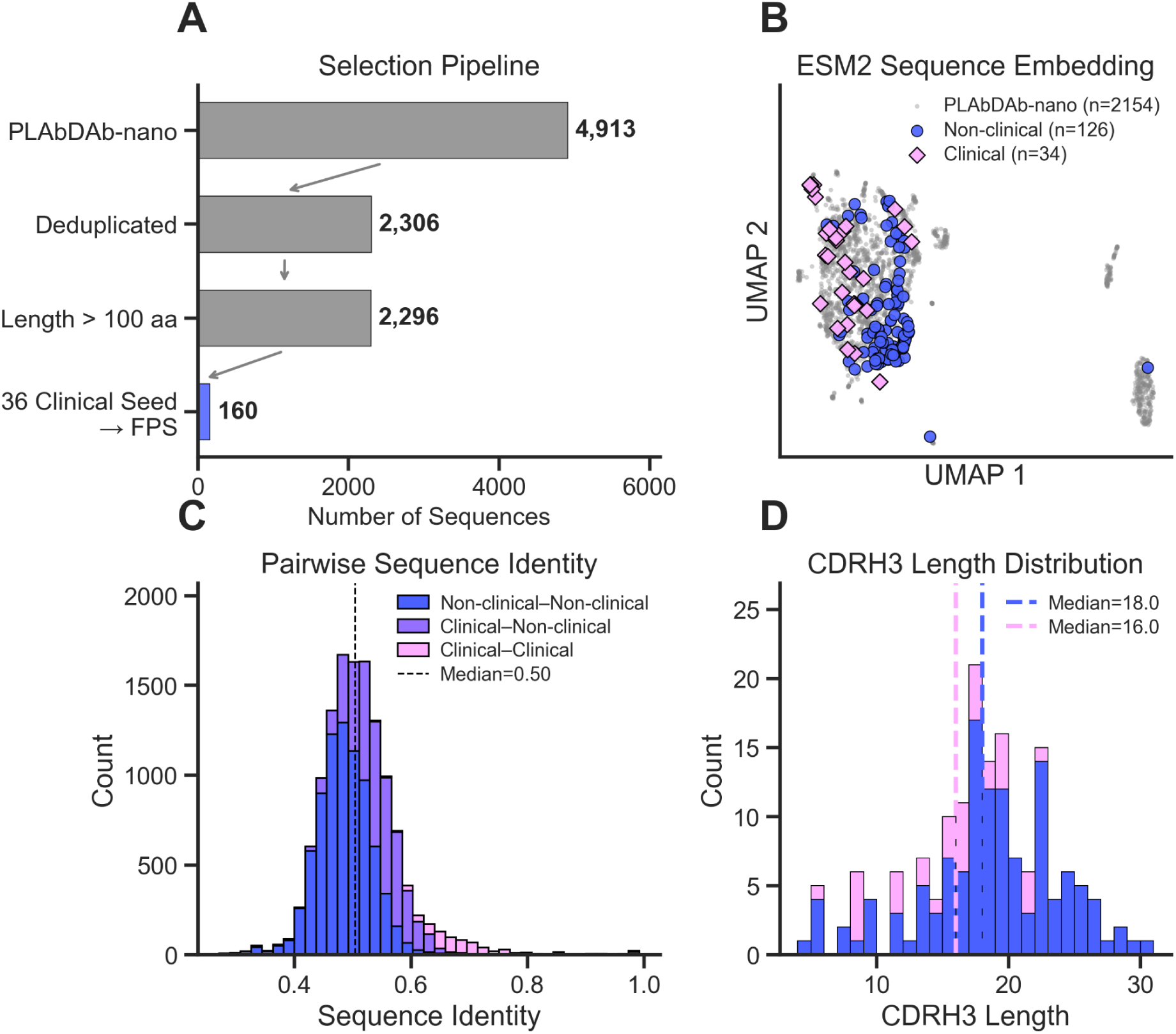
**Sequence diversity of the GDPa5 VHH-Fc library**. **A**. Selection of 160 VHH sequences from PLAbDAb-nano using farthest-point sampling seeded with 36 clinical nanobody sequences defined by Gordon et al. (2026), yielding a library designed to maximize sequence diversity while maintaining clinical relevance. **B**. UMAP projection of ESM2 sequence embeddings showing the distribution of GDPa5 library sequences relative to the broader PLAbDAb-nano sequence space and the clinical seed sequences. **C**. Pairwise sequence identity distributions for comparisons among clinical and non-clinical GDPa5 sequences, illustrating the broad sequence diversity of the selected library. **D**. CDRH3 length distributions for clinical and non-clinical GDPa5 sequences, demonstrating broader CDRH3 diversity among the non-clinical library members.

A UMAP projection of ESM2 sequence embeddings [Lin et al., 2023] confirmed that the GDPa5 library broadly spans both clinical and general single-domain antibody sequence space (**Figure 2B**). Pairwise sequence identity analysis colored by pair type, revealed a median identity of 52%, confirming that GDPa5 provides a diverse evaluation set (**Figure 2C**). CDRH3 lengths vary more for the non-clinical sequences (median 19.0, range 4-30) than in the clinical sequences (median 16, range 5-23), reflecting the broader sequence diversity of the PLAbDAb-nano pool (**Figure 2D**).

### VHH-Fc biophysical properties form orthogonal clusters that recapitulate IgG behavior

We determined the developability properties of the GDPa5 library using our reported biophysical assay panel, PROPHET-Ab [Arsiwala et al., 2025]. Hierarchical clustering of the assay results revealed three main clusters (**Figure 3A–B**). Self-association (AC-SINS) in two buffer conditions, namely PBS (pH 7.4) and 20 mM Histidine/20 mM NaCl (pH 6.0), clustered strongly (Spearman ρ = 0.73). Surface hydrophobicity and colloidal stability (HIC and SMAC) clustered reasonably well (ρ = 0.54), consistent with both assays measuring adsorption to hydrophobic matrices. Polyreactivity (PR) measures separated into two orthogonal sub-clusters: the two PR BVP blocking buffer conditions (SuperBlock and BSA) correlated tightly with each other (ρ = 0.85), whereas PR CHO and PR OVA formed a distinct cluster (ρ = 0.62). The divergence between these two polyreactivity sub-clusters (Ward distance = 1.41) likely reflects the different molecular surfaces presented by baculovirus particles versus soluble proteins.

**Figure 3:**
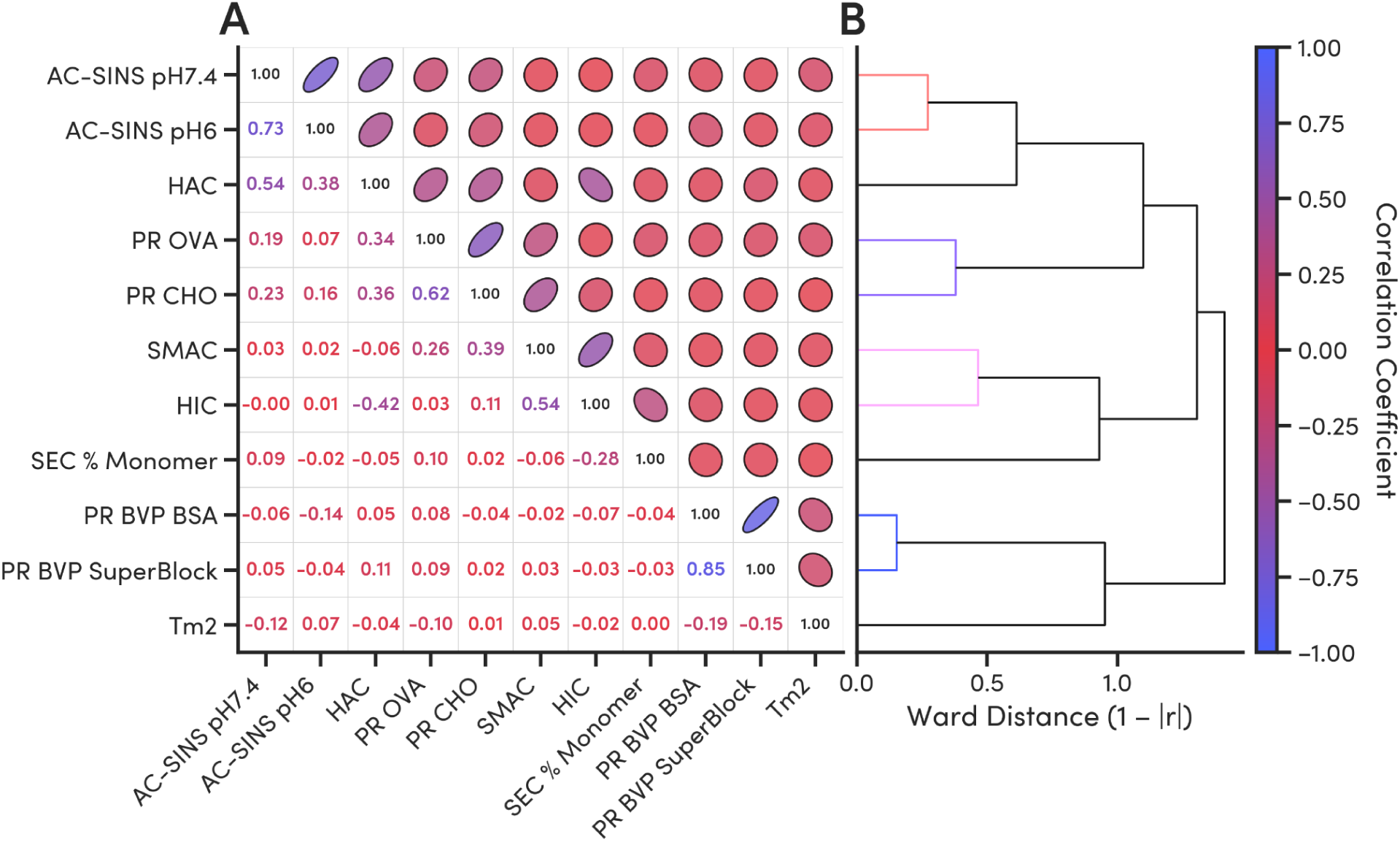
Relationships among biophysical developability properties of the GDPa5 VHH-Fc library. Analyses include the 100 VHH-Fcs with ≥80% monomer content by SEC. **A.** Spearman correlation matrix across developability assays. The lower triangle reports pairwise correlation coefficients, while the upper triangle provides an elliptical representation of the same correlations, with ellipse shape reflecting correlation magnitude and orientation reflecting direction. **B.** Hierarchical clustering of pairwise assay correlations using Ward distance, revealing distinct groupings of related biophysical properties, including self-association, hydrophobicity, and polyreactivity measurements.

Additionally, HAC showed modest positive correlations with PR OVA (ρ = 0.34) and PR CHO (ρ = 0.36), and a modest negative correlation with HIC (ρ = −0.42), consistent with electrostatic and hydrophobic surface patches representing partially competing structural features [Kraft et al., 2020]. Thermostability, as measured by the midpoint temperature of the second unfolding transition (Tm2), showed little correlation with any surface or interaction property, demonstrating that cooperative domain unfolding is mechanistically independent of surface-driven stickiness.

This clustering pattern closely mirrors the property interdependencies observed in the GDPa1 IgG dataset [Arsiwala et al., 2025], where HIC and SMAC, AC-SINS, and the two polyreactivity reagents formed analogous groupings. That VHH-Fcs recapitulate the same organization of biophysical properties as full-length IgGs suggests that the underlying physicochemical drivers of developability are conserved across these formats. We quantitatively demonstrate (**Figure S1)** that both the assay correlation matrices and the hierarchical assay clustering are correlated between GDPa5 and GDPa1 datasets with statistical significance.

**Figure S1.**
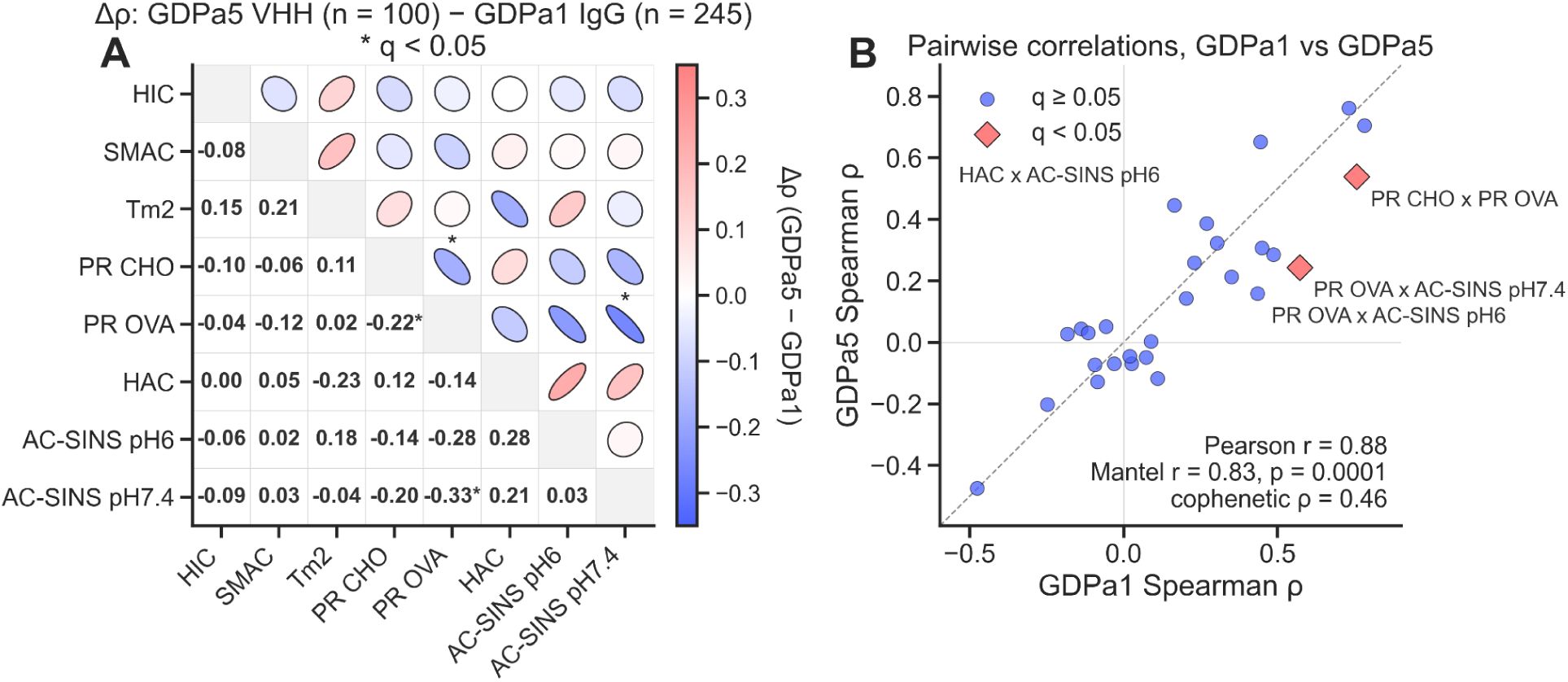
**Correlation of assays between GDPa5 and GDPa1**. **A.** Correlation matrix of differences between the correlation matrices between GDPa1 and GDPa5. Ellipses are colored by which dataset has a higher correlation between assays. Red is when GDPa5 is more correlated and blue is when GDPa1 is more correlated. Ellipse eccentricity is related to the degree of correlation. The q value is a threshold based on the Fisher-z test with the Benjamini-Hochman correction for correlation of independent Spearman correlations. A q < 0.05 represents a rejection of the null of correlation between the panels **B.** Correlation of Spearmans between GDPa5 and GDPa1 datasets.

### Core surface and stability assays validate robustly against an orthogonal dataset

Results for the GDPa5 library across the full PROPHET-Ab assay panel are shown in **Figure 4A–D**. Data are only shown for sequences that passed an 80% monomer threshold as determined by SEC, and based on the assay distribution **(Figure S2)**. Based on the distribution of SEC monomer content, an 80% monomer threshold was used to restrict subsequent analyses to predominantly monomeric VHH-Fc preparations. Library sequences (blue) and clinical seed sequences (pink) [Gordon et al., 2026] show broadly similar distributions for surface hydrophobicity (HIC, SMAC), heparin binding (HAC), and BVP-based polyreactivity. In other properties, however, clinical sequences tend toward higher self-association (AC-SINS; **Figure 4A**), and non-clinical sequences tend toward higher CHO- and OVA-based polyreactivity (**Figure 4C**) and lower thermostability (**Figure 4D**). These shifts are mostly consistent with the selection and optimization pressures that VHH therapeutics undergo during clinical development, which likely enrich for lower polyreactivity and higher thermal stability [Svenilov et al., 2023]. However, the shift towards higher self-association demonstrates the complex interplay between these properties for developable antibodies.

**Figure 4:**
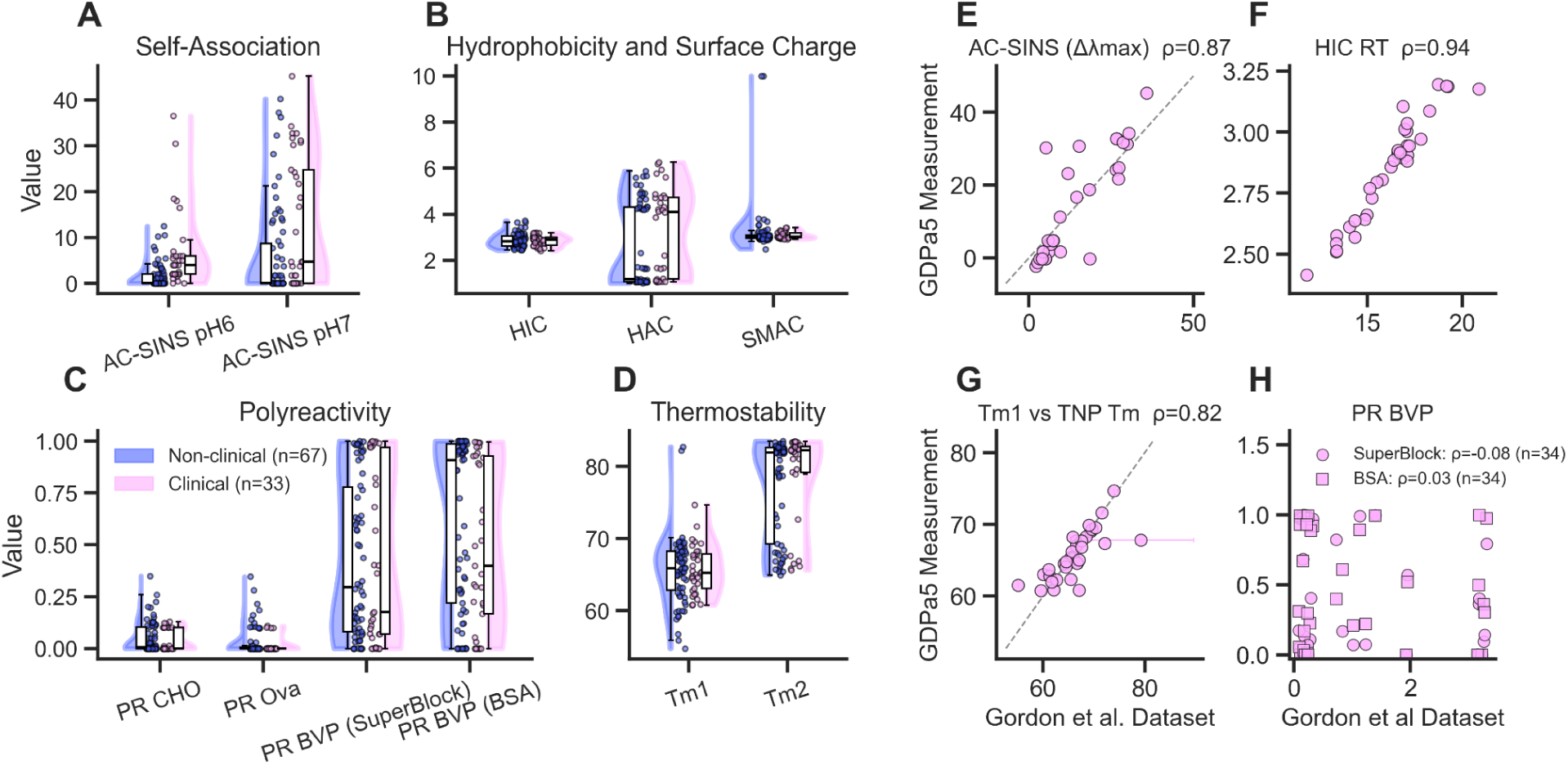
Distribution and cross-dataset comparison of VHH-Fc developability properties. Assay distributions are shown for GDPa5 VHH-Fcs with ≥80% monomer content by SEC. Clinical sequences overlapping with Gordon et al. 2026 are shown in pink, and non-clinical GDPa5 sequences are shown in blue. **A.** Self-association measured by AC-SINS in His/NaCl buffer at pH 6.0 and PBS at pH 7.4. **B.** Chromatographic measurements of hydrophobicity, heparin binding, and colloidal behavior by HIC, HAC, and SMAC, respectively. **C.** Polyreactivity measured against CHO cell lysate, ovalbumin, and baculovirus particles (BVP); BVP measurements were performed using SuperBlock and BSA blocking conditions. **D.** Thermostability determined from the second melting transition (Tm2), corresponding to unfolding of the VHH domain. For sequences shared between GDPa5 and Gordon et al., corresponding measurements were compared directly where matched assays were available. **E.** Cross-dataset comparison of AC-SINS measurements in PBS. **F.** Cross-dataset comparison of HIC retention times. **G.** Cross-dataset comparison of thermostability measurements (Tm1). **H.** Cross-dataset comparison of BVP polyreactivity measured under SuperBlock and BSA blocking conditions. Identity lines are shown for directly comparable measurements where appropriate. Error bars represent technical replicates for GDPa5 and reported variability from Gordon et al.; when not visible, error bars are smaller than the plotted symbols.

To further demonstrate the difference between the scaffolds, we look at the difference between the developability thresholds. Traditionally, the characterization for sufficient developability has been chosen by a 90% threshold of developability properties from clinical antibodies [Jain et al., 2017]. These thresholds were recently re-derived for a larger set of clinical antibodies. [Arsiwala et al., 2025]. For GDPa5, we take the clinical antibodies and derive another threshold using the same 90% value. The developability thresholds derived from the GDPa5 dataset differ from those established for clinical IgGs (**Figure S3**), reinforcing that IgG-derived cutoffs should not be applied directly to VHH-Fcs without format-specific recalibration.

**Figure S2.**
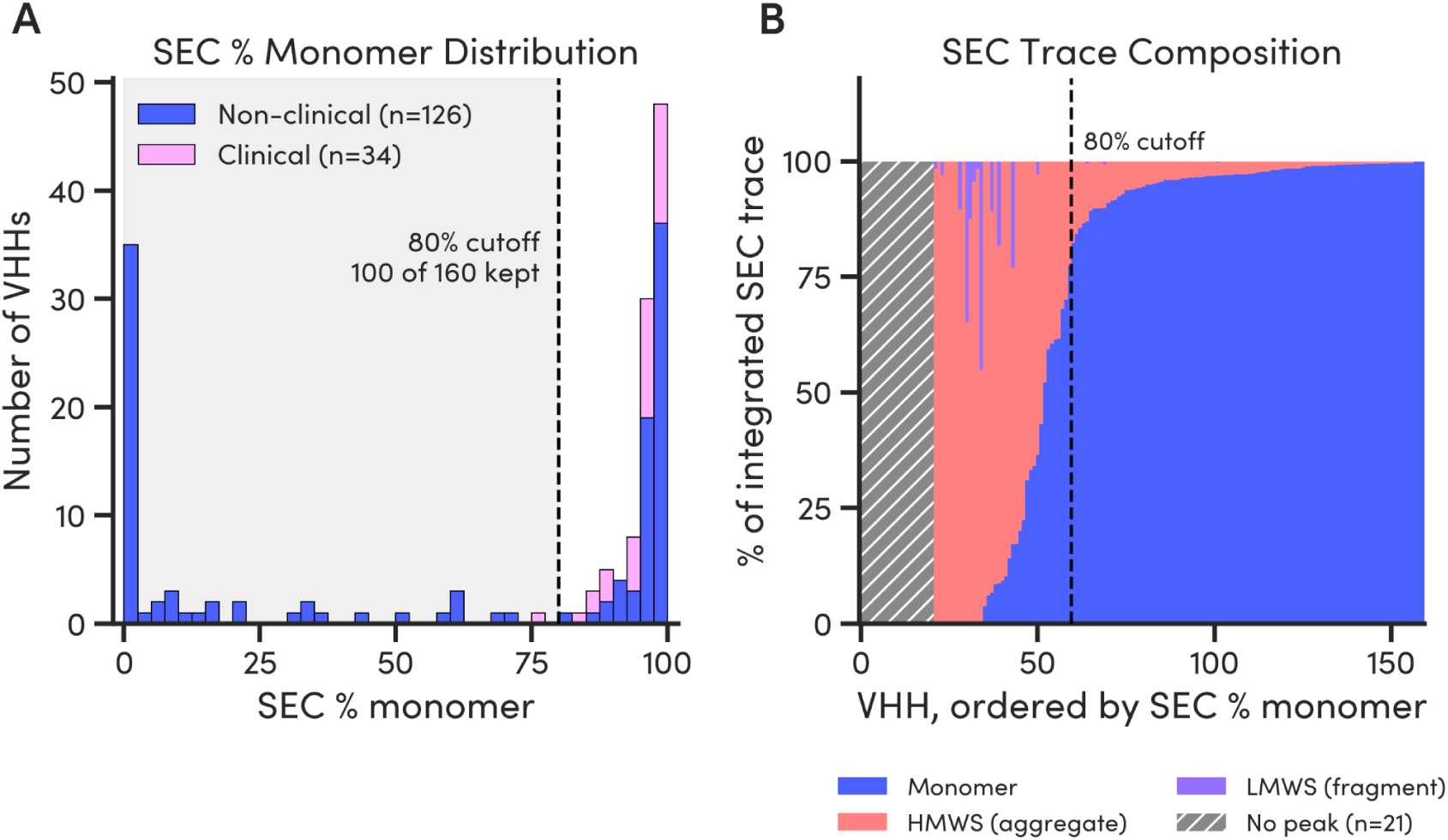
SEC monomer distribution and selection of the purity threshold for GDPa5 VHH-Fcs. **A.** Distribution of SEC monomer content across the GDPa5 library, with clinical sequences shown in pink and non-clinical sequences shown in blue. The 80% monomer threshold used for subsequent analyses is indicated. **B.** SEC composition of individual VHH-Fc preparations, ordered by decreasing monomer fraction, illustrating the relative contributions of monomeric and non-monomeric species across the library.

For the 34 sequences common to both GDPa5 and the Gordon et al. dataset, we assessed rank-order agreement between these independent measurements of the same biophysical properties. Surface and stability properties agree well: HIC (ρ = 0.94, **Figure 4F**), AC-SINS (ρ = 0.87, **Figure 4E**), and Tm1 (ρ = 0.82, **Figure 4G**). By way of contrast, polyreactivity (BVP) did not agree (ρ = −0.08 for SuperBlock blocking; ρ = 0.03 for BSA blocking, **Figure 4H**). We are unable to explain this discordance, and we encourage independent replication of this comparison to resolve it.

**Figure S3.**
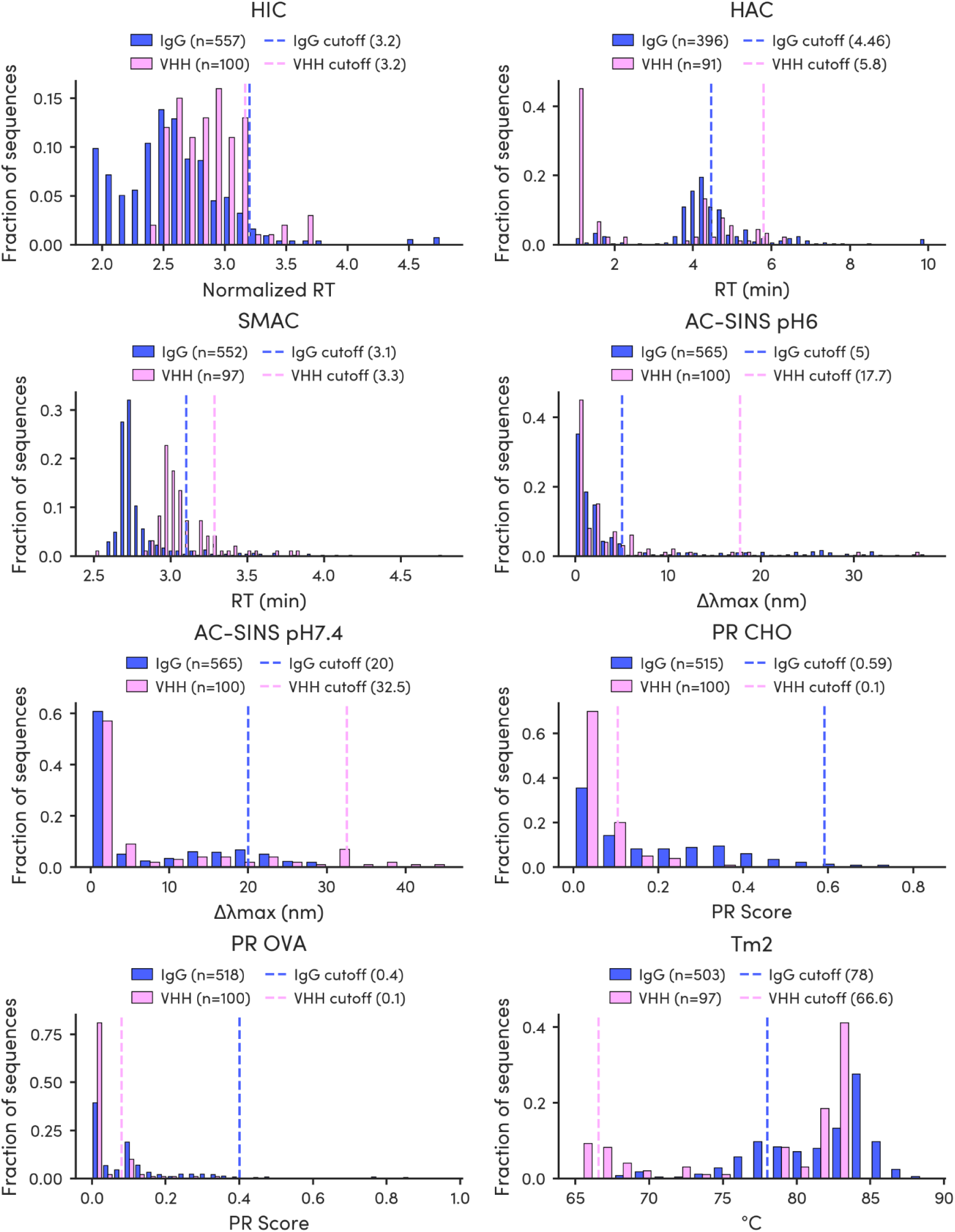
Cutoff thresholds comparing the clinical dataset 90% thresholds for IgGs and the 90% thresholds for the clinical VHH-Fcs in this dataset

### Intra-format VHH-Fc models are limited by data scarcity

To establish a baseline for intra-format prediction, we trained models exclusively on the GDPa5 VHH-Fc dataset using leave-one-cluster-out cross-validation (LOCO CV), where sequences were clustered by sequence identity to prevent data leakage and reflect the out-of-distribution challenge of real discovery campaigns (**Figure 5A**). To accommodate varying levels of model complexity, the following four architectures were considered: TabICLv2 [de Freitas et al., 2025], TabPFN v2.5 [Hollmann et al., 2023; Hollmann et al., 2025; Liu & Ye, 2025], XGBoost, and ridge regression. Both tabular neural networks, TabPFN v2.5 and TabICLv2, were considered using both fine-tuning (FT) and zero-shot (ZS). To represent the antibodies we evaluated featurizations including antibody-specific protein language model embeddings (AbLang2), general protein language model embeddings (ESM2), spatial structural features (DeepSP), isoelectric point (pI), and one-hot encodings (OHE). Ridge regression with OHE served as the null hypothesis baseline.

**Figure 5:**
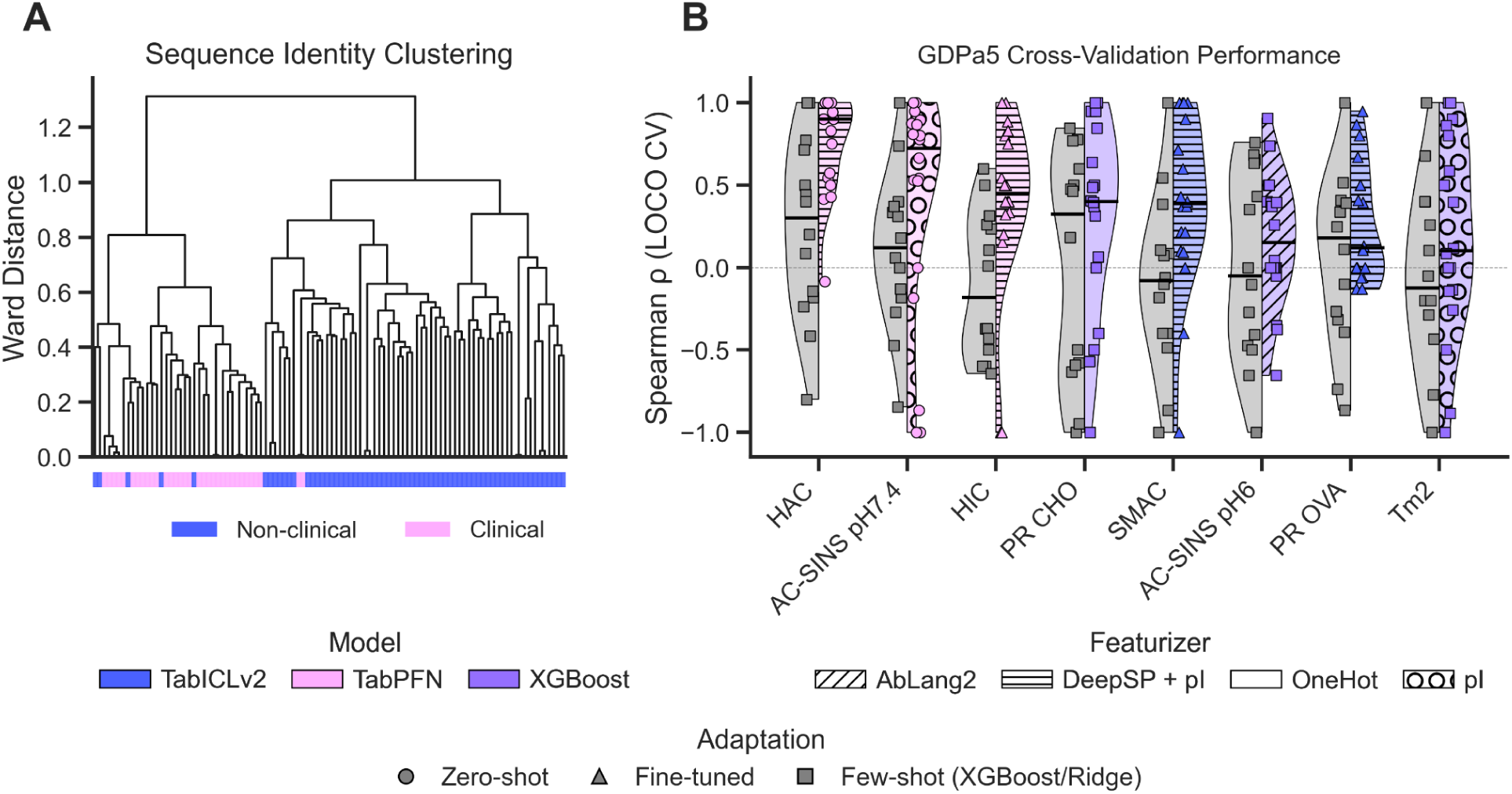
Model cross-validation performance on VHH-Fc sequence prediction. **A.** Hierarchical clustering representation of sequence similarity showing the clustering. Singleton clusters were removed from the evaluation. The colors below represent whether each sequence is a clinical or library sequence. **B.** Model performance using LOCO cross-validation. For each assay a baseline ridge regression (left) trained on one-hot encodings, and the best performing model (right) are shown. Colors represent the model.

Tabular models with richer featurization consistently outperformed the baseline, but absolute predictive performance was constrained across the panel (**Table 1**, **Figure 5B**). HAC retention time achieved the highest intra-format performance (ρ = 0.827). Notably, pI alone explained most of this signal (Ridge + pI: ρ = 0.795), consistent with heparin binding being driven primarily by surface charge. AC-SINS pH 7.4 (ρ = 0.607) and SMAC (ρ = 0.682) followed, while polyreactivity and thermostability proved most intractable (PR CHO ρ = 0.302, PR OVA ρ = 0.265, Tm2 ρ = 0.112). Ridge regression baselines were at or near zero for most assays, confirming that simple sequence encodings require extra information to be predictive. Critically, performance varied substantially across LOCO folds, indicating overfitting to individual sequence clusters rather than learning generalizable biophysical rules (**Figure 5B**). Scatter plots of pooled held-out predictions further illustrate this limited fidelity (**Figure S4**). Together, these results establish a clear ceiling on format-specific VHH-Fc prediction at N=100 and motivate the use of larger, cross-format IgG training data in the sections that follow.

**Figure S4:**
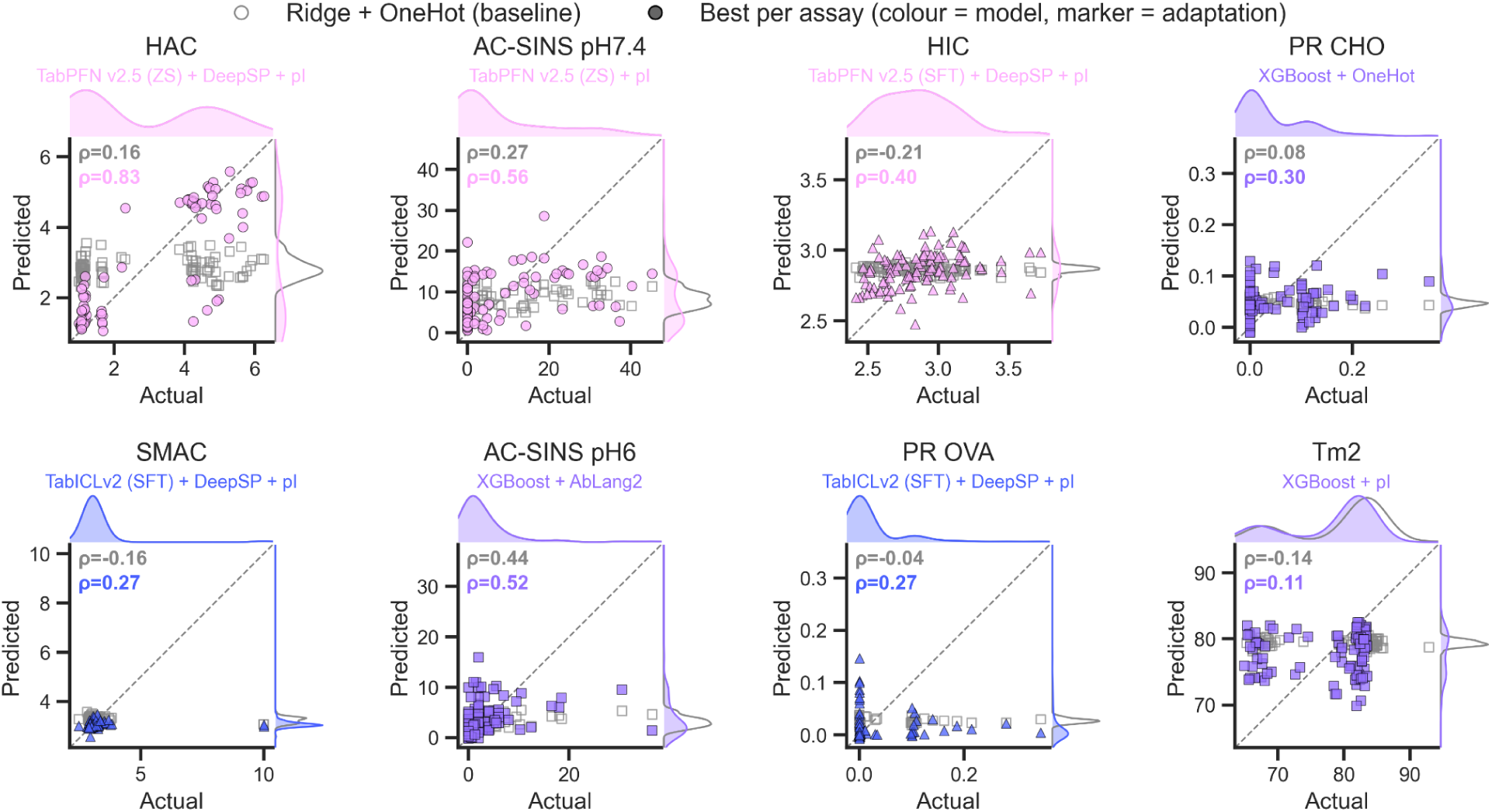
Scatter plot representations of the model performance on evaluation comparing the best model to the one-hot ridge baseline. Marginals correspond to the density of both distributions.

**Table 1:**
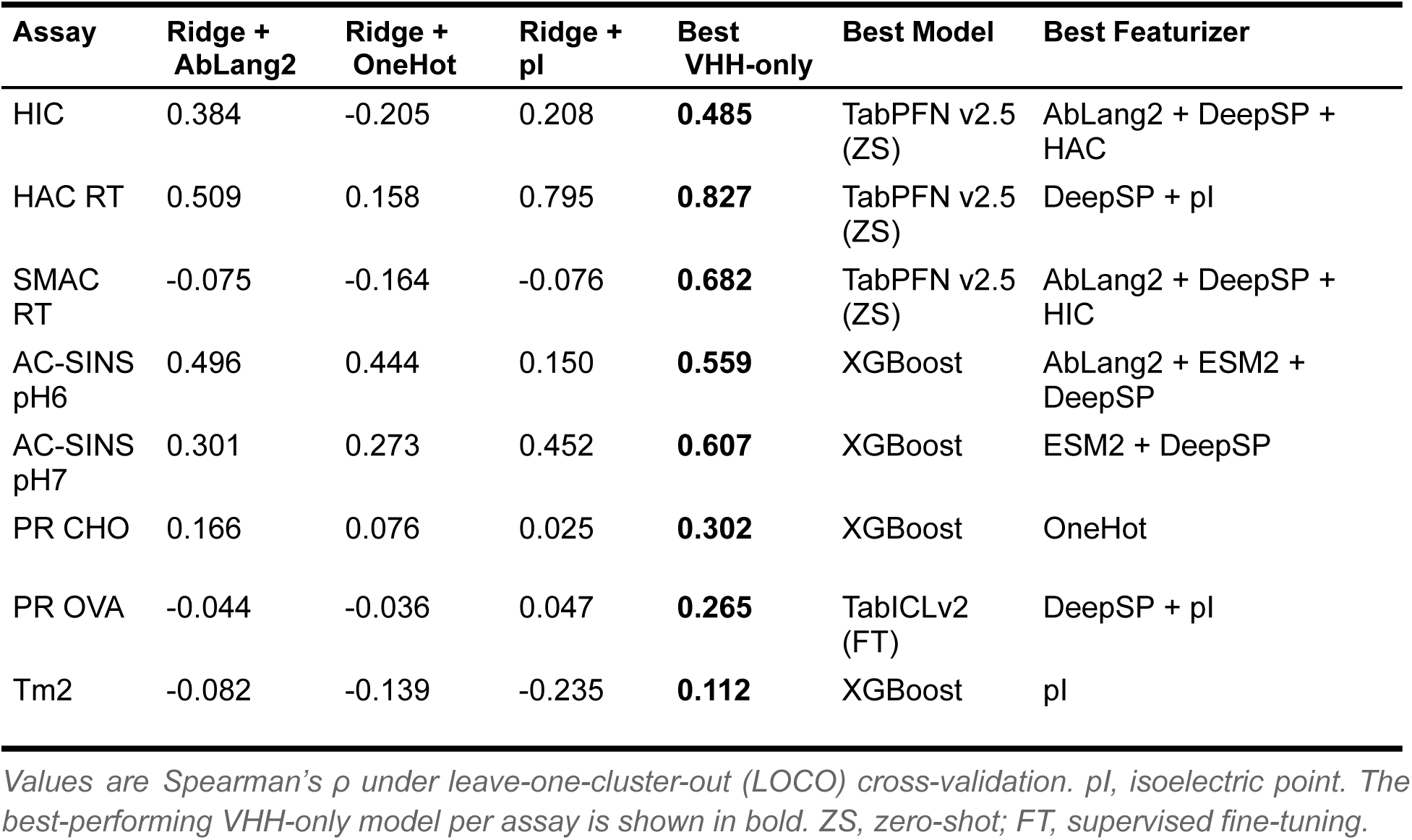
VHH LOCO model performance and comparison to baselines.

### VHH-Fc and IgG developability profiles differ while preserving shared biophysical relationships

Prior to cross-format modeling, we characterized the distributional shift between the 100 GDPa5 VHH-Fcs and the 559 standard IgGs that pass the same SEC threshold constituting our training set (**Figure 6**). The IgG training dataset comprises public datasets GDPa1 (n=246) [Arsiwala et al., 2025] and GDPa3 (n=80) [van Niekerk et al., 2026] and additional proprietary Ginkgo internal data (n=288). Sequence similarity analysis confirms that VHH variable domains are structurally distinct from IgG VH domains, with even the nearest IgG neighbor sharing limited sequence identity (**Figure 6A**). Violin plots of assay readouts reveal shifted baseline biophysical behaviors across formats. Polyreactivity (PR CHO, PR OVA), thermostability, and HAC retention times exhibit the most pronounced differences in both median and variance (**Figure 6B–E**). These shifts are mechanistically interpretable: VHH domains have evolved hydrophilic substitutions at residue positions corresponding to the VH-VL interface of conventional antibodies, compensating for the absent light chain by presenting a more hydrophilic surface [Muyldermans, 2013]. This altered surface character directly influences charge distribution and nonspecific binding propensity, shifting the biophysical baseline relative to IgGs. PR BVP assay data were not generated for the IgG training set and are therefore excluded from all cross-format modeling.

**Figure 6:**
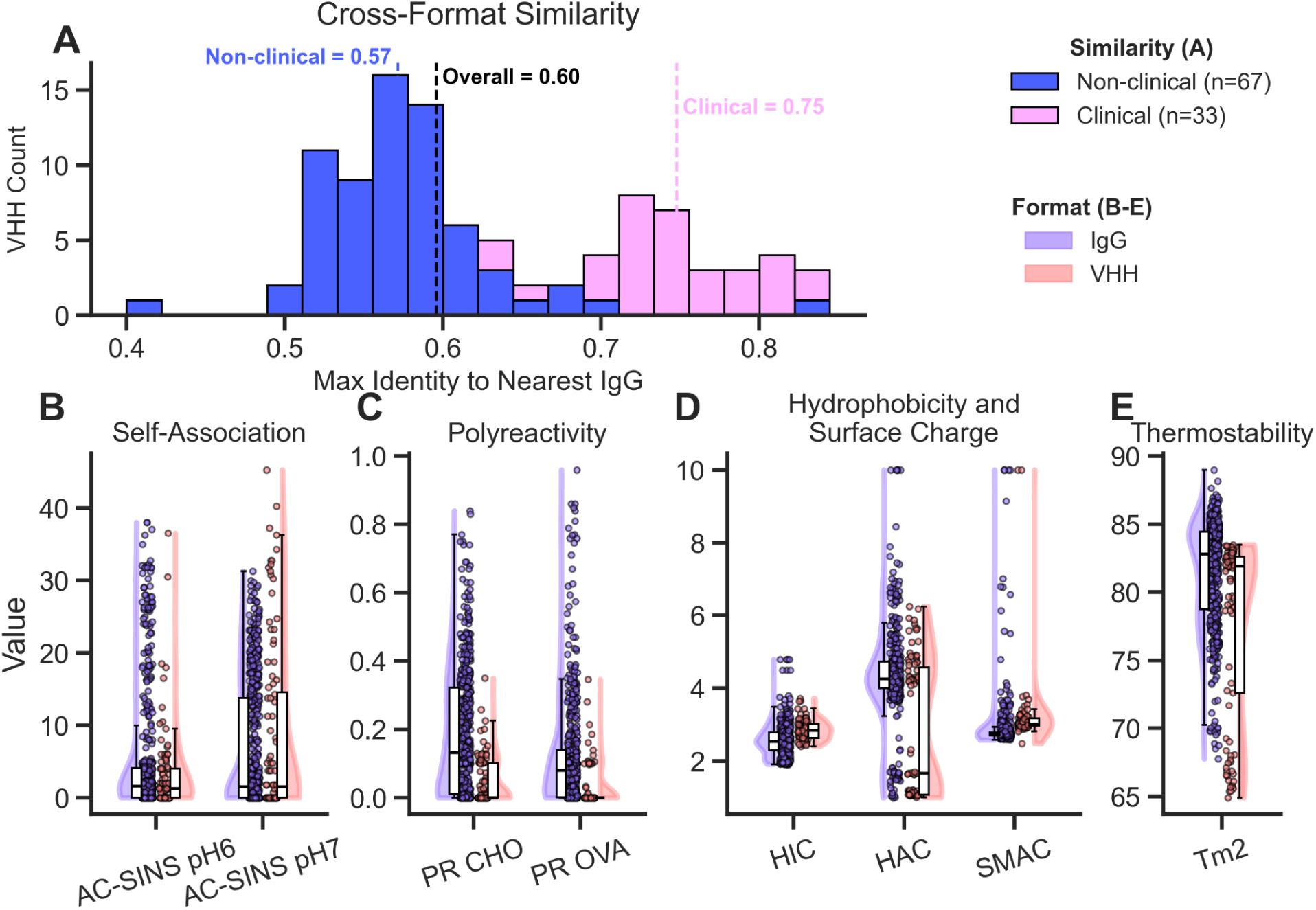
**Cross-format assay similarities**. These are only evaluated for the 100 VHH-Fcs with sufficient purity Violin plots representing the assay distributions for the respective datasets. **A.** Sequence similarity of the VHH sequences to the nearest IgG. The axes are grouped by assay type. **B.** Self-association via AC-SINS. **C.** Polyreactivity via PR CHO and PR OVA. **D.** Hydrophobicity and surface charge assays including HIC, HAC, and SMAC. **E.** Thermostability

Despite these distributional differences, the orthogonal property cluster architecture observed in GDPa5 mirrors that of the GDPa1 IgG dataset. In particular, the same property groupings, governed by the same underlying physicochemical drivers, are present in both formats. The preservation of these biophysical relationships across formats, despite differences in their absolute distributions, supports the feasibility of cross-format prediction. We test this hypothesis directly in the sections that follow.

### IgG-trained models generalize to VHH-Fc surface and self-association properties

Having established that our VHH-Fc set is data-limited, we trained models exclusively on the IgG heavy chains from the larger 559-IgG dataset and evaluated them on the GDPa5 VHH-Fc library - without any format-specific fine-tuning. For each assay, we performed a comprehensive ablation across all model architectures and featurization strategies. A full table of model performance for each ablation is available in the supplementary information. The best result per assay is reported in **Table 2** and **Figure 7A**. Tabular neural networks (TabICLv2, TabPFN v2.5) consistently outperformed ridge regression baselines across all assays, confirming that these architectures handle the distributional shift between IgG training data and VHH-Fc test data more effectively than conventional approaches. Scatter plots of the best model predictions are shown in **Figure 7B**.

**Figure 7:**
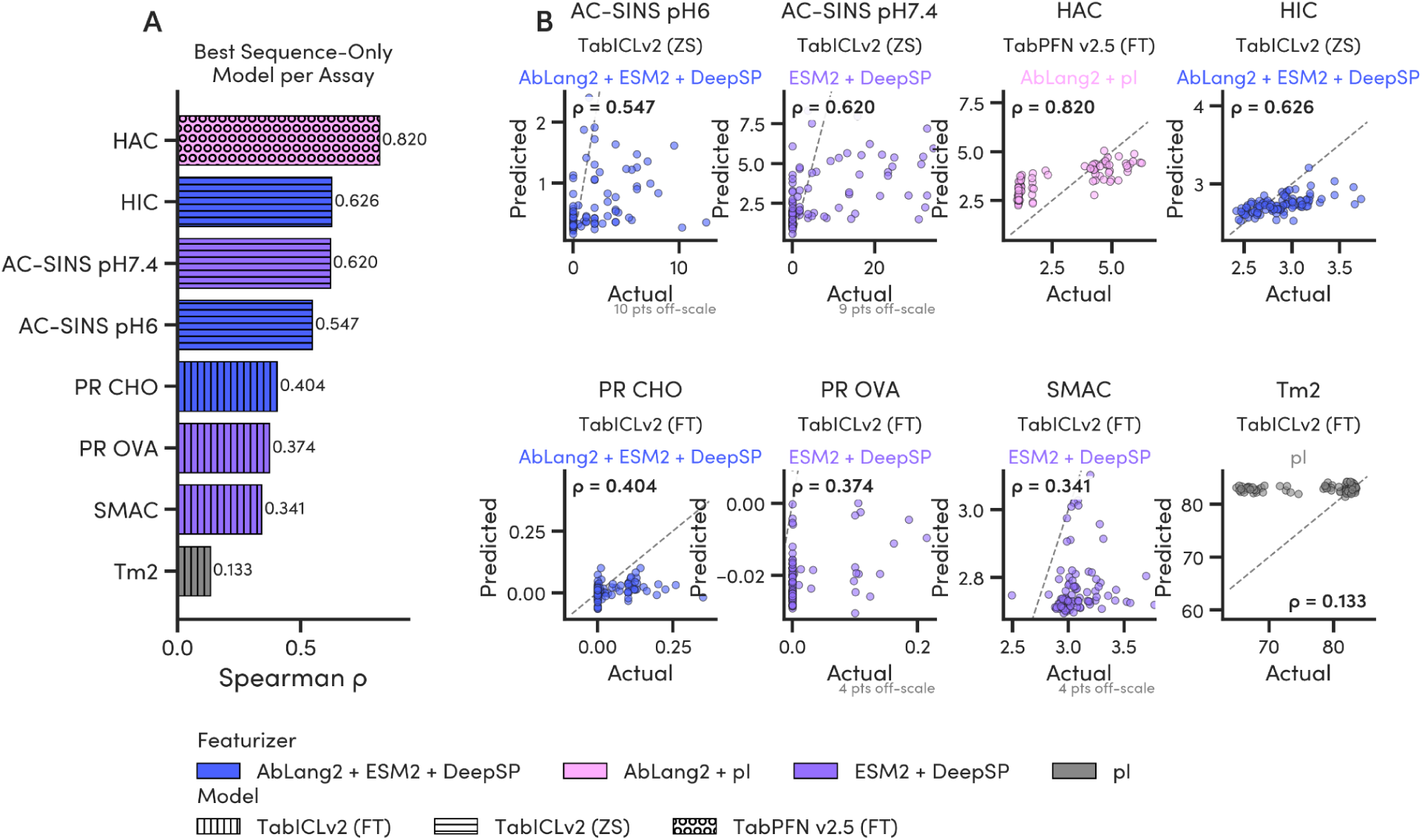
**Cross-format predictive model performance between IgG and VHH-Fcs**. For each model, the training data incorporated featurizations of the heavy chains from the 559 IgGs and the experimental assay results for the full mAb. The models were tested on the VHH-Fcs. **A.** The best performance model for each assay sorted in descending order by Spearman correlation coefficient. The bars are colored by featurization strategy and the hashes represent the model and training strategy. **B.** The corresponding scatterplot predictions on the test VHH-Fc set for the best models. Scatterplots are zoomed to the region of interest if the predictions do not follow the provided x=y line.

**Table 2:** Cross-format (IgG -> VHH-Fc) model performance and comparison to baselines.

| Assay | Ridge + AbLang2 | Ridge + OneHot | Ridge + pl | Ridge + best featurizer | Best sequence -only | Best + Auxiliary | Best Model | Best Featurizer |
| --- | --- | --- | --- | --- | --- | --- | --- | --- |
| HIC | -0.015 | -0.076 | -0.009 | 0.346 | **0.626** | 0.600 | TabICLv2 (ZS) | AbLang2 + ESM2 + DeepSP |
| HAC RT | 0.238 | 0.013 | 0.631 | 0.631 | 0.820 | **0.844** | TabPFN v2.5 (FT) | AbLang2 + pl + PR |
| SMAC RT | -0.048 | -0.136 | -0.081 | 0.059 | 0.345 | **0.734** | TabPFN v2.5 (FT) | DeepSP + HIC |
| AC-SIN S pH6 | -0.173 | 0.440 | 0.451 | 0.451 | 0.547 | **0.581** | TabICLv2 (FT) | AbLang2 + ESM2 + DeepSP + PR |
| AC-SIN S pH7 | 0.353 | 0.289 | 0.606 | 0.606 | 0.620 | **0.647** | TabPFN v2.5 (ZS) | ESM2 + DeepSP + HIC + HAC |
| PR CHO | 0.217 | 0.183 | 0.248 | 0.285 | 0.404 | **0.506** | TabICLv2 (ZS) | AbLang2 + pl + AC-SINS |
| PR OVA | 0.023 | -0.097 | 0.170 | 0.268 | 0.374 | **0.425** | TabPFN v2.5 (ZS) | ESM2 + DeepSP + AC-SINS |
| Tm2 | 0.110 | 0.155 | -0.009 | 0.155 | 0.155 | **0.187** | TabPFN v2.5 (ZS) | AbLang2 + DeepSP + PR |

Values are Spearman's $\rho$ for cross-format prediction (IgG-trained models evaluated on VHH-Fcs). pl, isoelectric point. The Ridge baselines use fixed featurizers; "Ridge + best featurizer" is Ridge with the best-performing sequence featurizer per assay. "Best sequence-only" uses sequence-derived features only; "Best + Auxiliary" adds measured or predicted assay inputs, and the higher of the two per assay is shown in bold. ZS, zero-shot; FT, supervised fine-tuning.

Three properties transferred with high cross-format fidelity. HAC retention time achieved the highest zero-shot performance (ρ = 0.820, TabPFN v2.5 FT, AbLang2 + pI), consistent with heparin binding being governed primarily by electrostatic contributions. HAC is a localized property that pI and antibody-specific language model embeddings capture well regardless of scaffold. Hydrophobicity (HIC) transferred similarly (ρ = 0.626, TabICLv2 ZS, AbLang2 + ESM2 + DeepSP), reflecting that exposed hydrophobic patches are a sequence-encoded feature conserved across formats. Self-association generalized reliably in both buffer conditions (AC-SINS pH 7.4, ρ = 0.620; AC-SINS pH 6.0, ρ = 0.547). Sequence-based predictions for polyreactivity (PR CHO ρ = 0.404; PR OVA ρ = 0.374) and colloidal stability (SMAC ρ = 0.345) were moderate. These properties depend on a more complex interplay of surface features and are addressed through the auxiliary cascade described in the next section.

Thermostability (Tm2) had near-zero zero-shot performance (ρ = 0.155). This failure is mechanistically interpretable: thermal unfolding in IgGs is driven in large part by the cooperativity of the CH1-CL interface, which is absent in VHH-Fcs. The Tm2 transition in a VHH-Fc reflects melting of the VHH domain alone, a structurally distinct process that IgG-trained models have limited ability to predict. Thermostability therefore remains a format-specific property requiring direct experimental measurement. Cross-format performance for transferable properties improved monotonically with IgG training set size (**Figure S5**), reinforcing that the scale of the IgG training dataset is a primary driver of cross-format predictive fidelity.

### Augmenting IgG models with easily measured surface properties unlocks prediction of complex liabilities

Properties governed by discrete, localized surface features transfer cleanly in zero-shot prediction. Complex properties such as polyreactivity and colloidal stability, however, arise from a dynamic interplay of hydrophobic patches, surface charge, and conformational flexibility that sequence embeddings alone cannot fully resolve. We reasoned that surface properties already predicted with high fidelity, HIC, HAC, or AC-SINS, could serve as auxiliary inputs to improve prediction of these harder-to-predict properties. Critically, these auxiliary values can either be rapidly measured early in a screening campaign or substituted with the sequence-only cross-format predictions described in the previous section, making the cascade computationally self-contained. We tested this by augmenting cross-format tabular models with auxiliary assay features and report improvements where Δρ ≥ 0.05, reflecting the practical cost of incorporating additional experimental measurements (**Figure 8A**).

**Figure 8:**
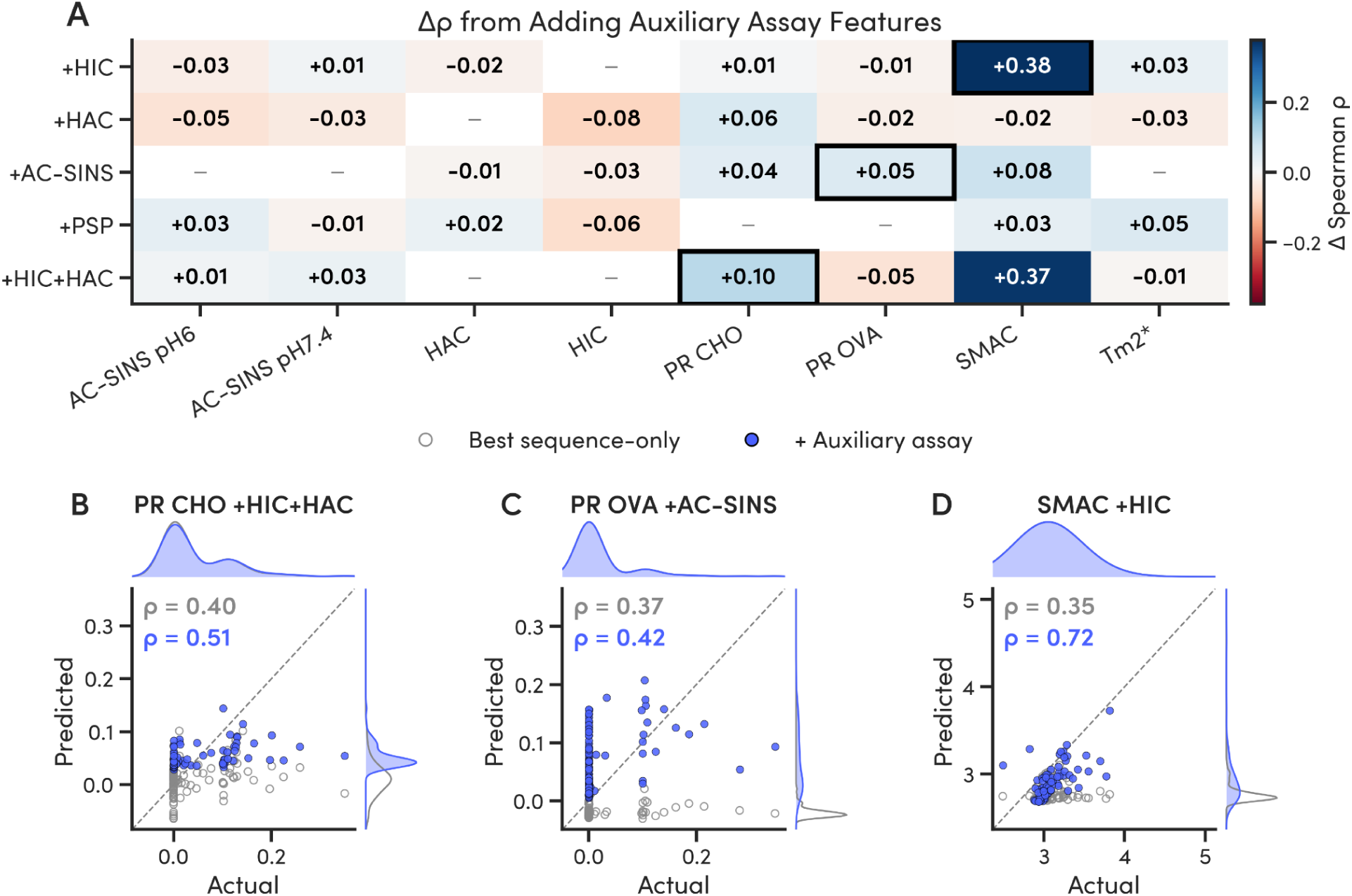
Auxiliary assay data for hydrophobicity and heparin binding improves cross-format performance of more complex assays, such as polyreactivity and SMAC. For Tm2, each tile corresponds to a different featurization strategy. **A.** A heatmap demonstrating the change in spearman correlation coefficient for each assay with the addition of auxiliary assay feature(s). The corresponding scatterplots of the best sequence-only model (grey) and auxiliary model predictions (blue) for **B.** SMAC + HIC, **C.** PSP CHO + HIC + HAC, and **D.** PSP Ova + AC-SINS.

The most informative gains came for polyreactivity, a complex, multi-modal liability that sequence embeddings alone predict only moderately. For polyreactivity to CHO cell lysate (PR CHO), incorporating experimental values for HIC and HAC together improved prediction from ρ = 0.404 to ρ = 0.506 (Δρ = +0.102, **Figure 8C**), consistent with polyreactivity arising from the combined effect of hydrophobic patches and positive surface charge. [Ritter et al., 2026]. Adding experimental AC-SINS to the PR OVA model yielded a more modest but meaningful improvement (ρ = 0.374 to ρ = 0.425, Δρ = +0.051, **Figure 8D**), suggesting a shared self-association component in ovalbumin-based polyreactivity. In both cases, the auxiliary inputs are distinct from the target property, so the gains reflect genuine cross-property information rather than a correlated readout of the same measurement. Colloidal stability (SMAC) showed the largest nominal gain from adding experimental HIC (ρ = 0.345 to 0.734, Δρ = +0.389, **Figure 8B**), but this improvement is expected and less informative [Walbi et al., 2021]. HIC and SMAC both report hydrophobic adsorption and are strongly correlated (**Figure 3**), so the experimental HIC value largely substitutes for a proxy measurement of the same property rather than resolving an independent liability. Thermostability (Tm2) remained essentially unpredictable even with auxiliary inputs (Δρ = +0.05), confirming that no combination of surface properties can substitute for scaffold-specific cooperative unfolding information. Full model-level results for each auxiliary combination are provided in **Figure S5**.

These findings establish a practical tiered triage cascade for VHH-Fc developability prediction. Surface hydrophobicity and heparin binding, which are the most reliably predicted properties, serve as inputs to polyreactivity - a complex liability that sequence alone predicts poorly. This hierarchical approach converts the cross-format prediction problem from a single hard task into a sequence of progressively informed predictions, substantially extending the practical utility of IgG training data for VHH-Fc developability screening. Additionally, this prediction cascade highlights a future path of which assays can be reliably replaced with *in silico* methods and which require more data to reach that point.

**Figure S5.**
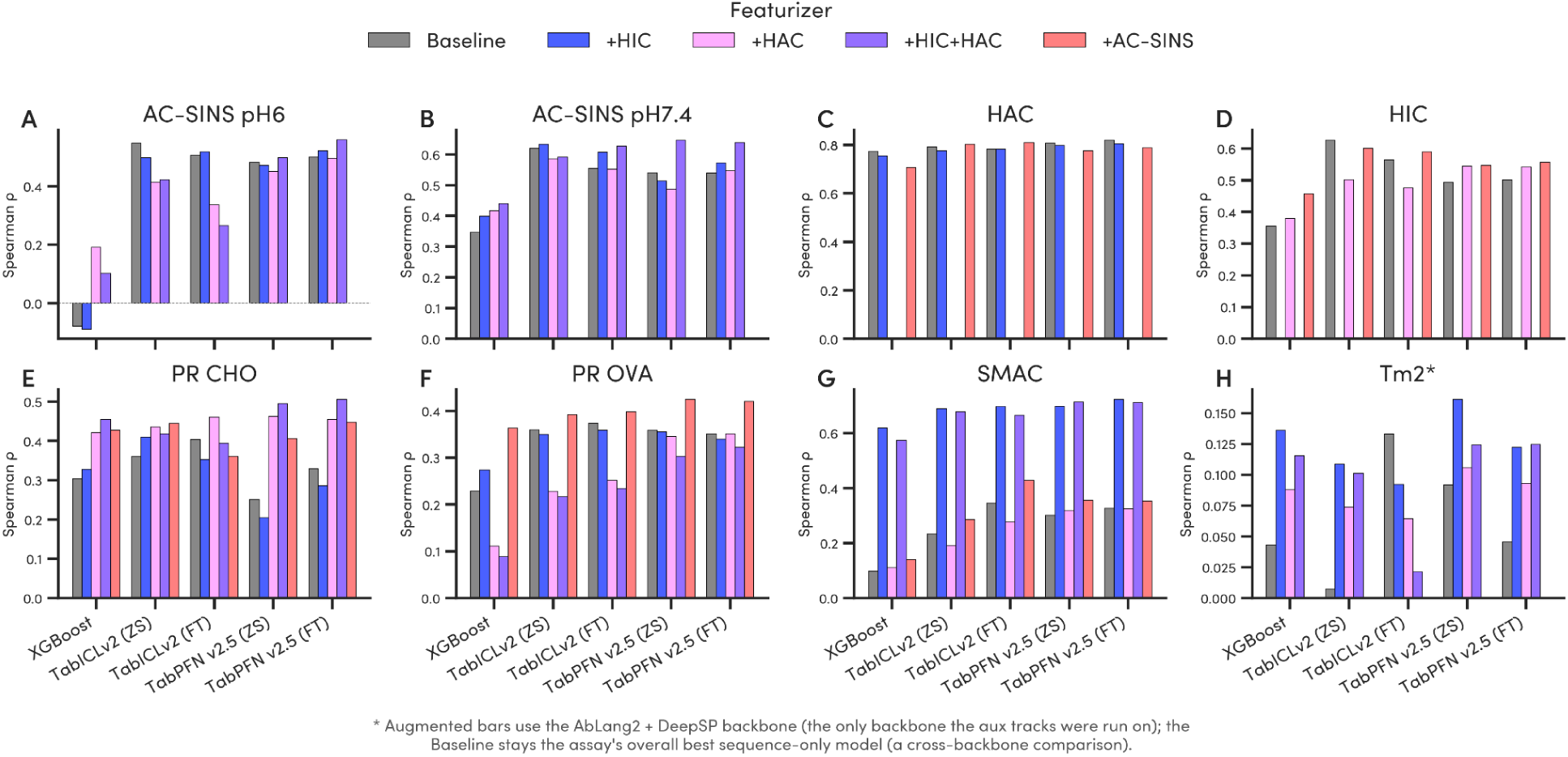
Improvement of auxiliary assays on prediction of properties that increased the most with experimental assay information as a function of assay and model

### Training-set scale enables IgG models to outperform VHH-Fc-specific approaches

To complete the evaluation, we directly compared the best IgG cross-format models against the best intra-format VHH-only LOCO CV models on each property (**Figure 9A**). This comparison is the critical test of the central premise. Despite its structural divergence from VHH-Fcs, IgG data provides more generalizable predictive signal than format-specific data at the scale available. The IgG-trained models outperformed VHH-only models across the panel in test conditions, often by substantial margins. HIC showed the largest gain (Δρ = +0.14; cross-format ρ = 0.626 vs. VHH-only ρ = 0.485). PR CHO and PR OVA showed the largest relative improvements (Δρ = +0.20 and +0.16 respectively), reflecting the near-complete failure of VHH-only models for polyreactivity at this sample size. Improvements for HAC (Δρ = +0.02), SMAC (Δρ = +0.05), and AC-SINS pH 6.0 (Δρ = +0.02) were more modest but consistent. For Tm2, both approaches remain effectively unpredictable, as established above. Scatter plots comparing the best cross-format (purple) and VHH-only (orange) model predictions confirm these differences visually across the full test set (**Figure 9B**).

**Figure 9.**
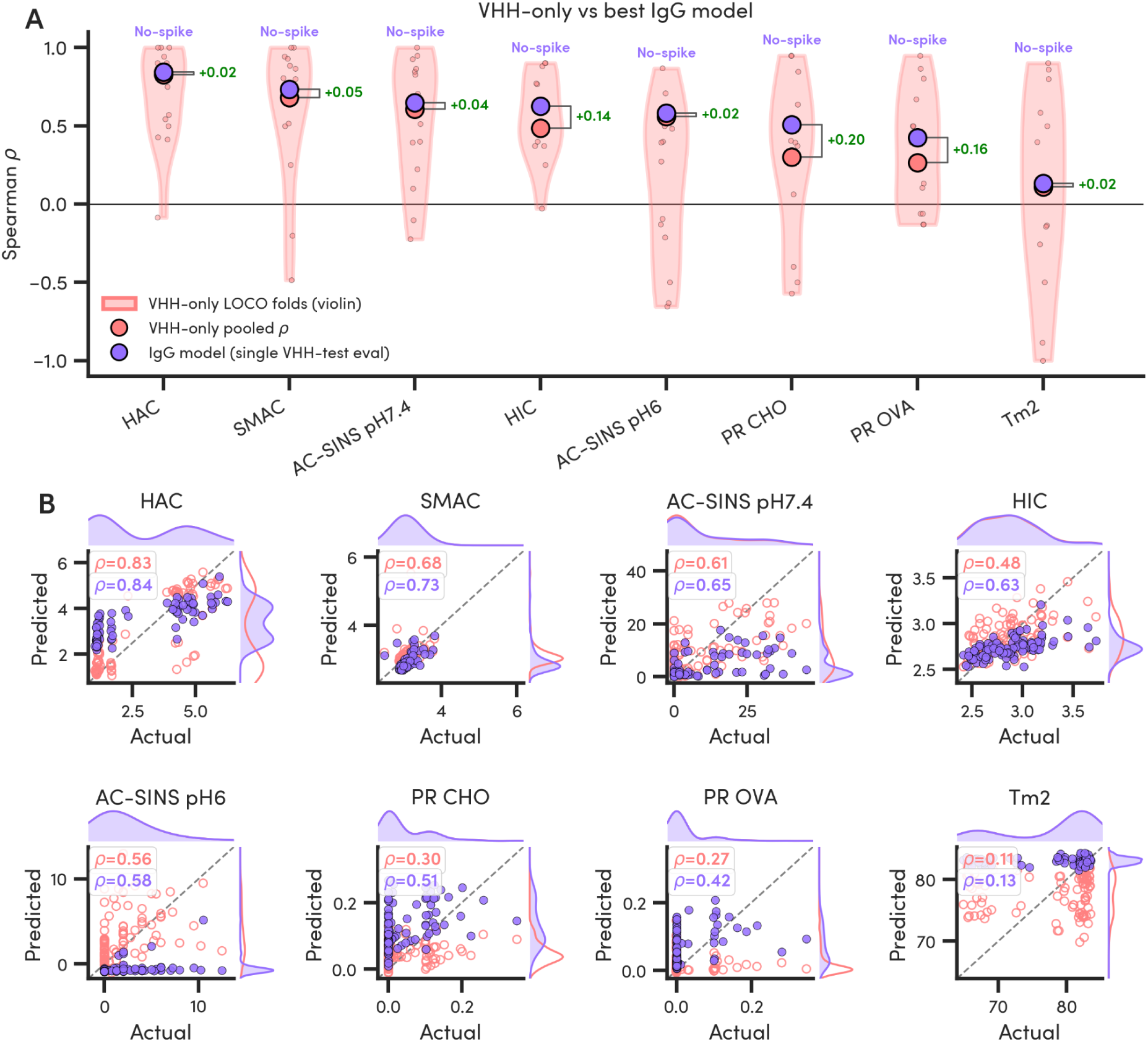
Overall comparison of cross-format prediction to VHH-trained models. **A.** Violin plot showing VHH-only LOCO CV folds with the pooled Spearman correlation coefficient as a large orange dot. The purple corresponds to the best IgG-based model for each property prediction on the VHH-Fcs data. The amount of spike-in data required to improve the model is shown above. The purple dot in that case corresponds to the pooled SCC for LOCO CV. **B.** Scatter plots of prediction of the best model for each property with cross-format (purple) and VHH-Fc only (orange). Densities of label distributions and predictions are shown on the marginals.

To test whether augmenting IgG training data with small amounts of VHH-Fc data could further improve performance, we evaluated a range of spike-in percentages (**Figure S6**). For all assays, incorporating VHH-Fc data did not substantially improve over the zero-shot IgG baseline. AC-SINS pH 7.4 was the single exception. A 25% spike-in yielded a minor improvement of Δρ = +0.02. This performance gain is not substantial enough for the spike-in gain and therefore is not used as the IgG-based reference for this property in **Figure 9A**. With this exception, the consistent zero-shot cross-format performance across multiple properties demonstrates that IgG data can provide a practical and scalable foundation for VHH-Fc developability prediction. We also show the change to these models’ predictive capability if we were to be more lenient on the SEC % threshold and demonstrate prediction almost entirely decreases (**Figure S7**). This is true for all properties except for PR CHO, but we suspect this is an artifact of the data distribution of these sequences. For a full breakdown of the contributions of each feature to the overall prediction power, we refer the reader to **Figure S8.** The sequence diversity and scale of the IgG dataset provide tabular neural networks with a broader landscape of sequence-property rules than can be extracted from 100 format-specific examples, enabling generalization across the structural divergence between IgGs and VHH-Fcs.

**Figure S6.**
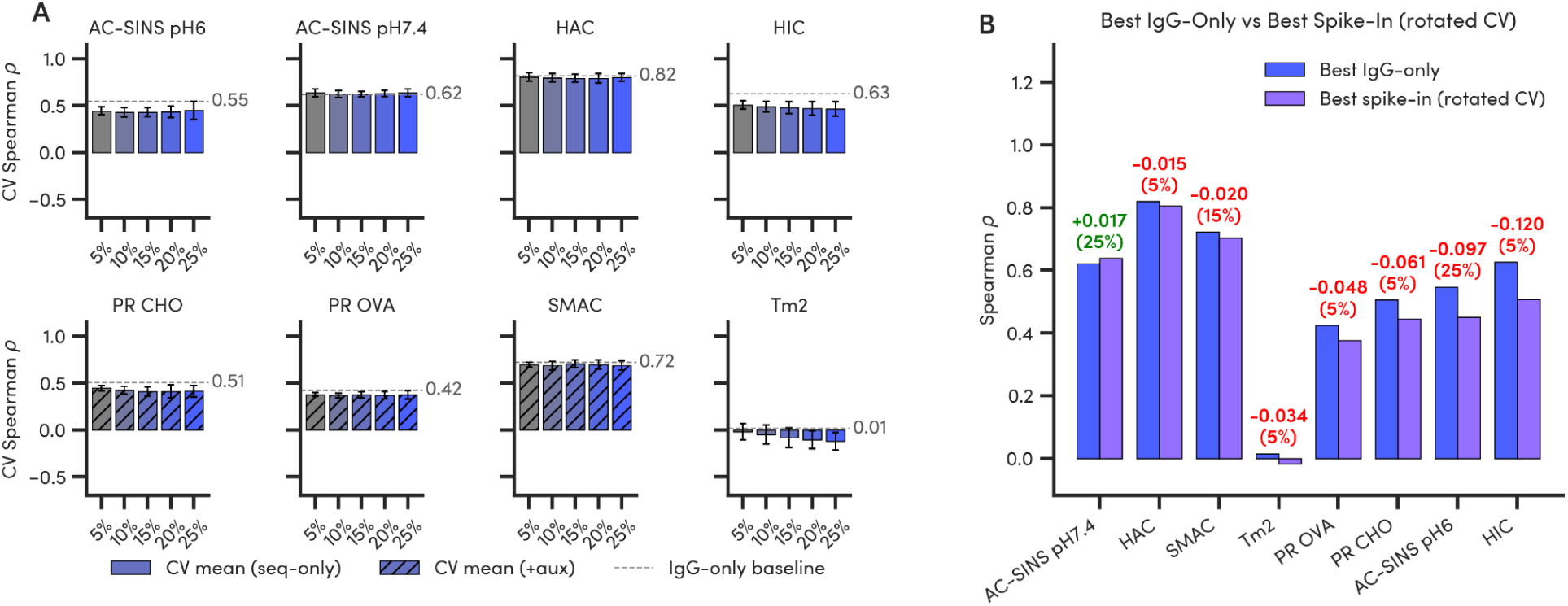
Cross-format model performance improvement when incorporating spike-in percentages of VHH-Fcs into the IgG-based models. **A.** Bargraphs with variances of each of the spike-in models when considering a LOCO CV strategy. Hashes correspond to those that performed best with an auxiliary property. **B.** Overall comparison of best spike-in to best model on each property

**Figure S7.**
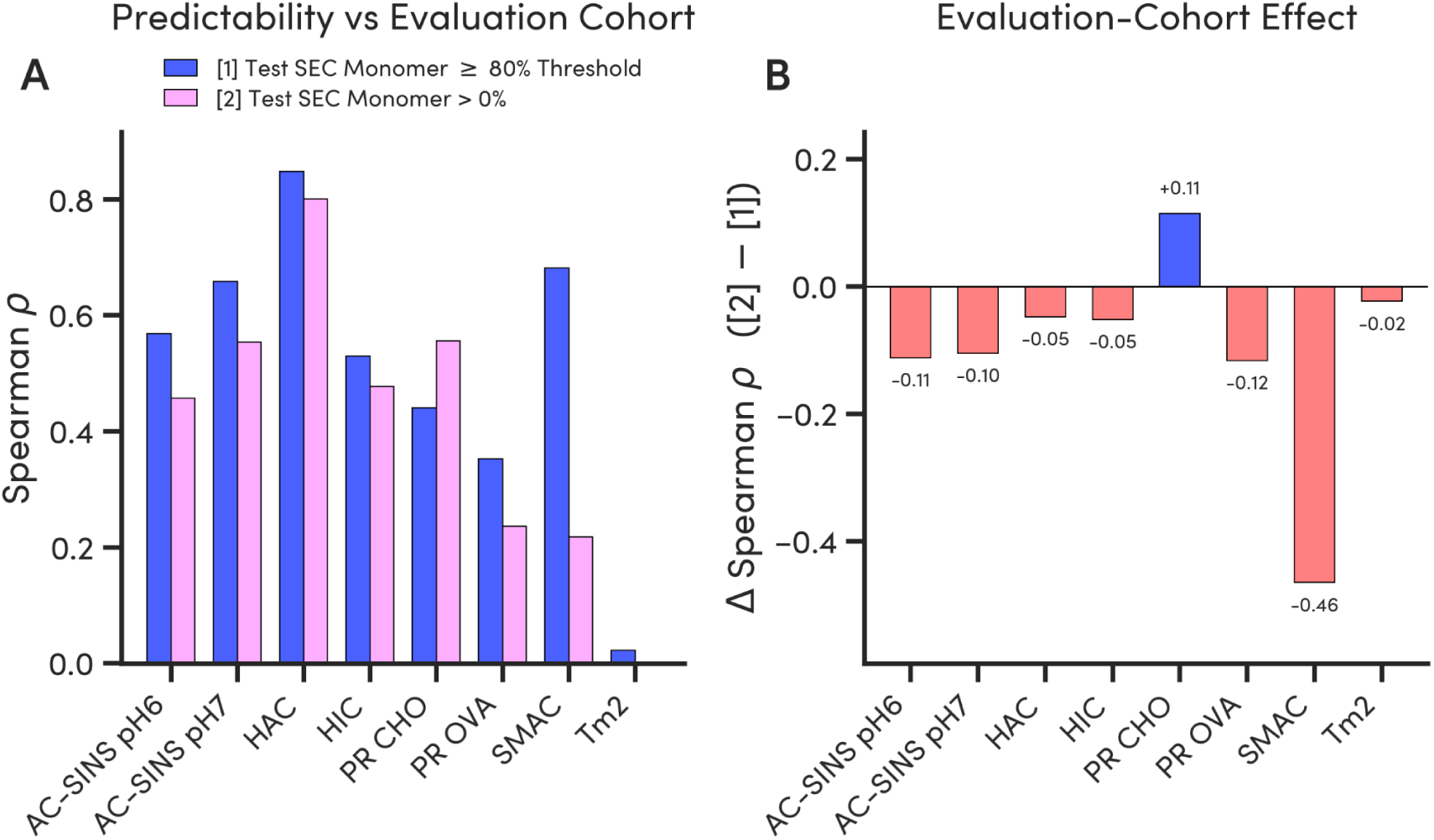
Model performance difference between SEC thresholds. **A.** Bargraphs demonstrating the difference between the best model for each assay testing on the full dataset of VHH-Fcs (n=126) where monomer threshold is above 0% (pink) and 80% (blue) **B.** Bargraph representing the difference in Spearman between 0% threshold arm and the 80% threshold arm.

**Figure S8.**
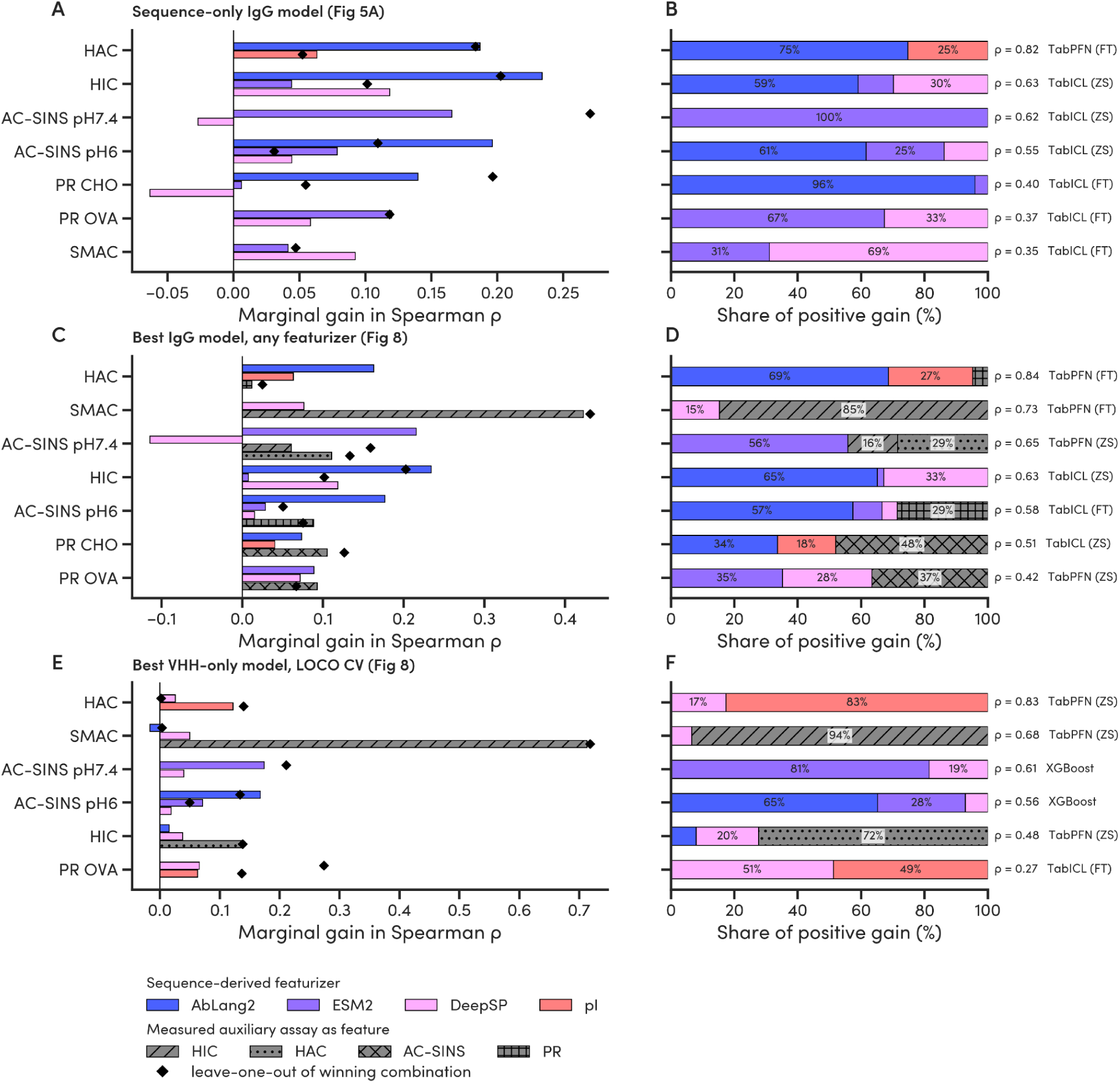
Featurizer-class contributions to the winning model of each modelling track. For each assay, the model with the highest held-out Spearman rho was decomposed by ablation over the featurizer combinations already evaluated for that same assay and model. Left column (**A, C, E**): marginal gain in rho from each featurizer class. Bars give the mean change over all observed pairs of runs whose featurizer sets differ only by that class and black diamonds show the effect of dropping the class from the winning combination itself, plotted where that run exists. Negative bars mean the class lowered rho on average. Right column (**B, D, F**): the same means as each class’s share of the total positive gain, with the winning model and its rho at the right. Tracks are the sequence-only IgG-to-VHH model (**A, B**), the best IgG-to-VHH model over all featurizers including measured auxiliary assays (**C, D**), and the best VHH-only model under LOCO CV (**E, F**). Colors mark sequence-derived featurizers, and hatched grey bars an auxiliary assay used as a feature. Single-class winners cannot be decomposed and are not included.

To determine how much IgG training data is required to surpass format-specific prediction, we retrained the best model for each assay on increasing subsets of the IgG dataset and evaluated cross-format performance on the VHH-Fc library (**Figure 10A**). As a control for model complexity, we applied Ridge regression and XGBoost using the same optimal featurization as the best model for each assay. The two simpler models behaved similarly to one another and scaled slowly with added data, most visibly for HAC, AC-SINS, and PR OVA. The best model was not always superior at the smallest training size, but it scaled more steeply and overtook the simpler models as data accumulated. This pattern was clearest for AC-SINS and HIC, indicating that the tabular neural networks capture cross-format signals that shallower models do not.

**Figure 10.**
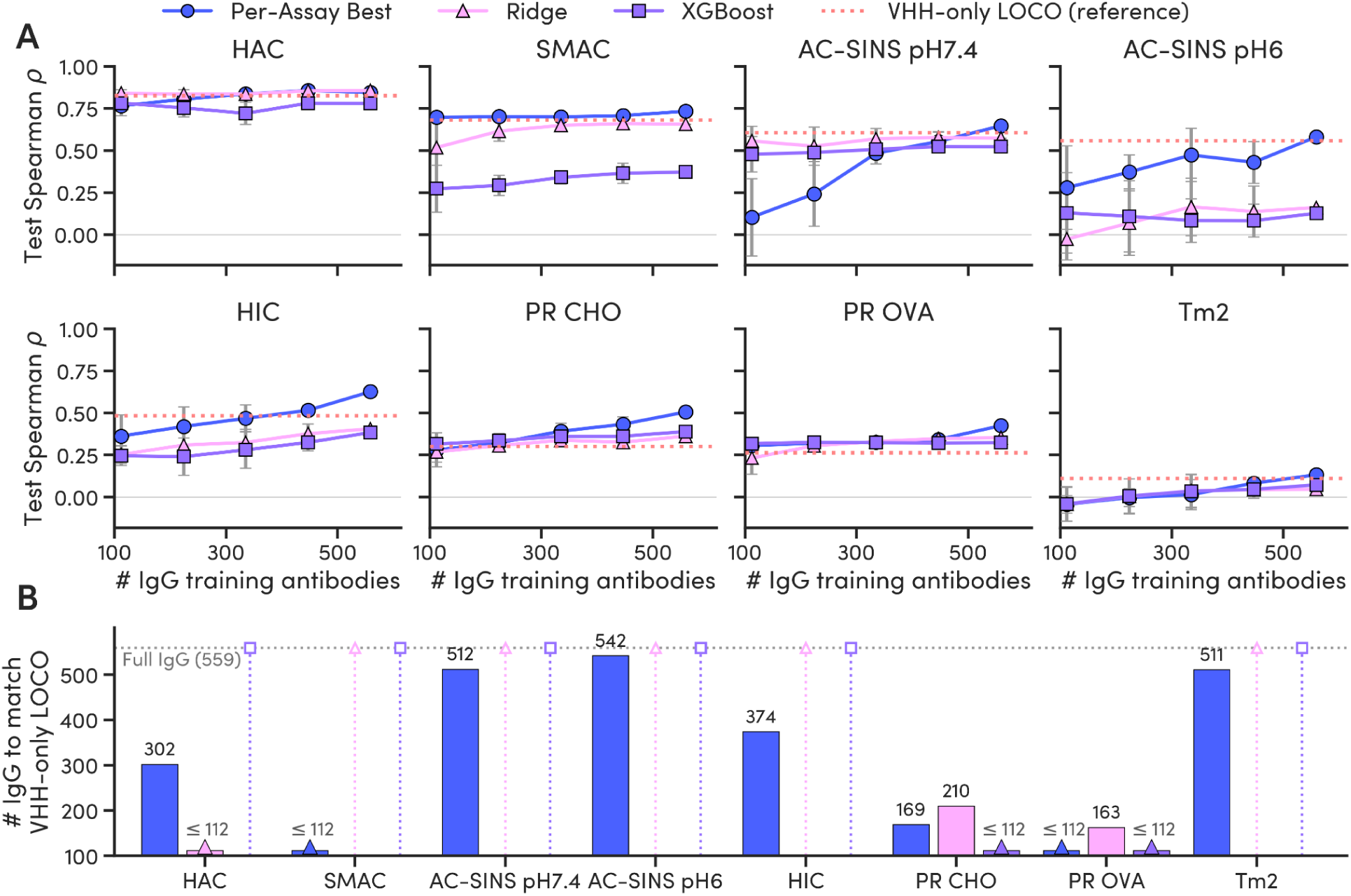
Cross-format performance scales with IgG training set size. For each assay, the best-performing cross-format model (Per-Assay Best) was retrained on increasing subsets of the IgG training data and evaluated on the VHH-Fc test set. Ridge regression and XGBoost, each using the same optimal featurization as the best model for that assay, are shown as controls for model complexity. **A.** Test Spearman ρ as a function of IgG training set size. The red dotted line marks the VHH-only LOCO baseline (Figure 4). **B.** The IgG training set size at which each model first exceeds the VHH-only baseline. Bars labeled “≤112” denote models that surpass the baseline at the smallest subset tested (the 20% point); bars reaching the dashed line (“Full IgG, 559”) denote models that never exceed it within the available data. Error bars in A represent the standard deviation across cross-validation folds.

We then interpolated the training set size at which each model first exceeds the VHH-only baseline (**Figure 10B**). The best model crossed this threshold for every assay, though the data required varied widely, from fewer than 112 antibodies to more than 500 (AC-SINS, Tm2). For HAC, PR CHO, and PR OVA, one of the models surpassed the baseline at low data scale through featurization rather than architecture. In these cases, Ridge regression or XGBoost with the optimal feature set performed comparably to the best model. The tabular neural networks were most advantageous for properties that require large training sets to transfer, such as AC-SINS and HIC, and less critical where a strong feature set alone sufficed.

## Discussion

This work tests directly whether format-specific training data is required for format-specific developability prediction, a question that had not been systematically evaluated. By profiling a diverse 160-member VHH-Fc library across 10 biophysical assays and evaluating cross-format prediction from a relatively larger 559-antibody IgG training set, we demonstrate that models trained entirely on IgGs systematically outperform models trained on VHH-Fcs alone. The explanation is not that VHH-Fc and IgG developability are identical (the distributional shifts in **Figure 5** make clear that they are not), but that the physicochemical rules governing surface-driven biophysical properties are conserved across the structural divergence between formats. Additionally, the sequence diversity and the size of the larger IgG dataset (n=559) provides a richer landscape for learning those rules than a 160 format-specific VHH dataset can.

The mechanistic basis for cross-format transfer follows from the nature of the properties that transfer successfully. Hydrophobicity, heparin binding, and self-association are governed by localized, surface-encoded features, such as exposed hydrophobic patches, charge distribution, and isoelectric point, that are sequence-determined and format-independent. These features are captured effectively by antibody-specific protein language models and structural descriptors regardless of the molecular scaffold. There is recent evidence supporting that the heavy chain surface characteristics primarily dictate these properties, further proving the possibility of this cross-format evaluation. [Sakhnini et al., 2026] Properties that do not transfer cleanly, such as thermostability and polyreactivity, require information about the full molecular context. Polyreactivity is partially rescued by the auxiliary cascade because its multi-modal surface drivers can be approximated from other measurable properties (HIC and HAC). Thermostability cannot be rescued because it reflects cooperative domain unfolding that is fundamentally different between IgGs and VHH-Fcs.

The failure of cross-format Tm2 prediction is potentially mechanistically revealing. Thermal unfolding in IgGs is dominated by cooperativity of the CH1-CL interface, which is absent in VHH-Fcs. The Tm2 transition in a VHH-Fc reports on the isolated VHH domain, which is a structurally distinct process for which IgG-trained models carry no relevant information. Taken with the GDPa4 findings [Ritter et al., 2026], where thermostability was similarly the least predictable property from parental IgG measurements across bispecific format transitions, a consistent pattern emerges: thermostability is a holistic, scaffold-dependent property that resists cross-format prediction regardless of the format transition. The convergence between GDPa4 and GDPa5 extends further. In both cases, surface hydrophobicity and heparin binding were the most reliably predicted properties and the most useful auxiliary features. This consistency across two structurally distinct format transitions suggests that the hierarchy of biophysical predictability identified in GDPa4 may be a general principle of the antibody developability landscape rather than a format-specific observation. This is also consistent with recent work understanding viscosity and pinocytosis across multispecific formats [Kraft et al., 2019; Fernandez-Quintero et al., 2023].

To our knowledge, this is the first demonstration of zero-shot, IgG-trained cross-format developability prediction for VHH-Fc antibodies. Prior work on cross-format transfer has been limited in scope, restricted to single properties or closely related format transitions [Xin et al., 2025]. This prior work has not tested the full developability panel across a structural gap as large as that between IgGs and VHH-Fcs. VHH-Fcs lack both the light chain and CH1 domain present in IgGs, yet cross-format prediction succeeds and does so better than format-specific models. This work shows that for properties governed by localized, sequence-encoded surface features, format-specific training data is not a prerequisite - provided sufficient cross format data is available. The practical implication of this work is immediate: groups working on any emerging antibody format lacking large-scale developability data can leverage existing IgG datasets as a productive starting point rather than waiting for format-specific experimental campaigns to accumulate.

To our knowledge, this is also the first application of tabular neural network foundation models to antibody developability prediction. Tabular neural networks have only recently been applied to protein property prediction [Guan et al., 2026]. Ridge regression baselines performed at or near zero for most cross-format prediction tasks, confirming that linear models do not handle the distributional shift between IgG training data and VHH-Fc test data. TabICLv2 and TabPFN v2.5 leverage in-context learning and meta-learned priors to smooth over heterogeneous feature spaces and generalize across distributional shifts, which are capabilities directly suited to the cross-format prediction challenge. This finding is particularly relevant in light of the 2025 Ginkgo Datapoints Antibody Developability Competition outcomes [van Niekerk et al., 2026], which demonstrated that standard models overfit to training distributions and generalize poorly to held-out test sets. Tabular foundation models represent a promising architectural response to this persistent bottleneck, and their strong performance here supports their broader adoption in developability modeling.

## Conclusions and Future Directions

This study makes three contributions to antibody developability modeling: (1) We release GDPa5, a diverse 160-member VHH-Fc developability dataset profiled across 10 assays on the PROPHET-Ab platform, the first large-scale, standardized public dataset of its kind for this format. (2) We demonstrate that IgG-trained tabular neural networks predict VHH-Fc developability better than VHH-specific models in zero-shot, with surface and self-association properties transferring with high fidelity (HAC ρ = 0.820, HIC ρ = 0.626, AC-SINS ρ = 0.620) and thermostability remaining a format-specific exception. (3) We establish a tiered auxiliary assay cascade that extends cross-format performance for polyreactivity (PR CHO Δρ = +0.102), a complex liability poorly predicted from sequence alone, by incorporating easily measured or reliably predicted surface properties (HIC, HAC) as inputs.

Several limitations of this study should be noted. The GDPa5 library comprises 100 sequences after purity filtering, limiting statistical power for intra-format analyses. All VHH-Fcs were profiled on a single Fc scaffold, leaving the influence of Fc variants uncharacterized. IgG training data uses full-mAb assay measurements with models trained on heavy chain sequences only, introducing a structural mismatch that likely attenuates performance for light-chain-influenced properties. The optimal balance of cross-format and format-specific training data across different sample sizes and properties remains open, as the AC-SINS pH 7.4 spike-in result illustrates.

Several directions will build on this foundation. As has been demonstrated in Grinsztajn et al. [Grinsztajn et al., 2025] the distribution and type of synthetic tasks used for tabular foundation model pre-training can dramatically impact utility on particular classes of problems. Designing synthetic tasks better reflective of the functional priors of protein properties broadly, and antibody developability properties specifically, is likely to yield higher-performing models. Incorporating VHH-specific structural descriptors, particularly those capturing CDR3 loop conformations and the hydrophilic framework substitutions at the former VH-VL interface, could reduce the remaining gap for polyreactivity prediction from sequence alone. As IgG datasets grow beyond the current 559-antibody scale, cross-format performance for transferable properties should continue to improve, as the scaling analysis in **Figure S9** suggests. Extending this cross-format framework to other emerging formats, such as scFvs, Fabs, minibinders, and multispecific architectures, will test the generality of the principles established here.

**Figure S9.**
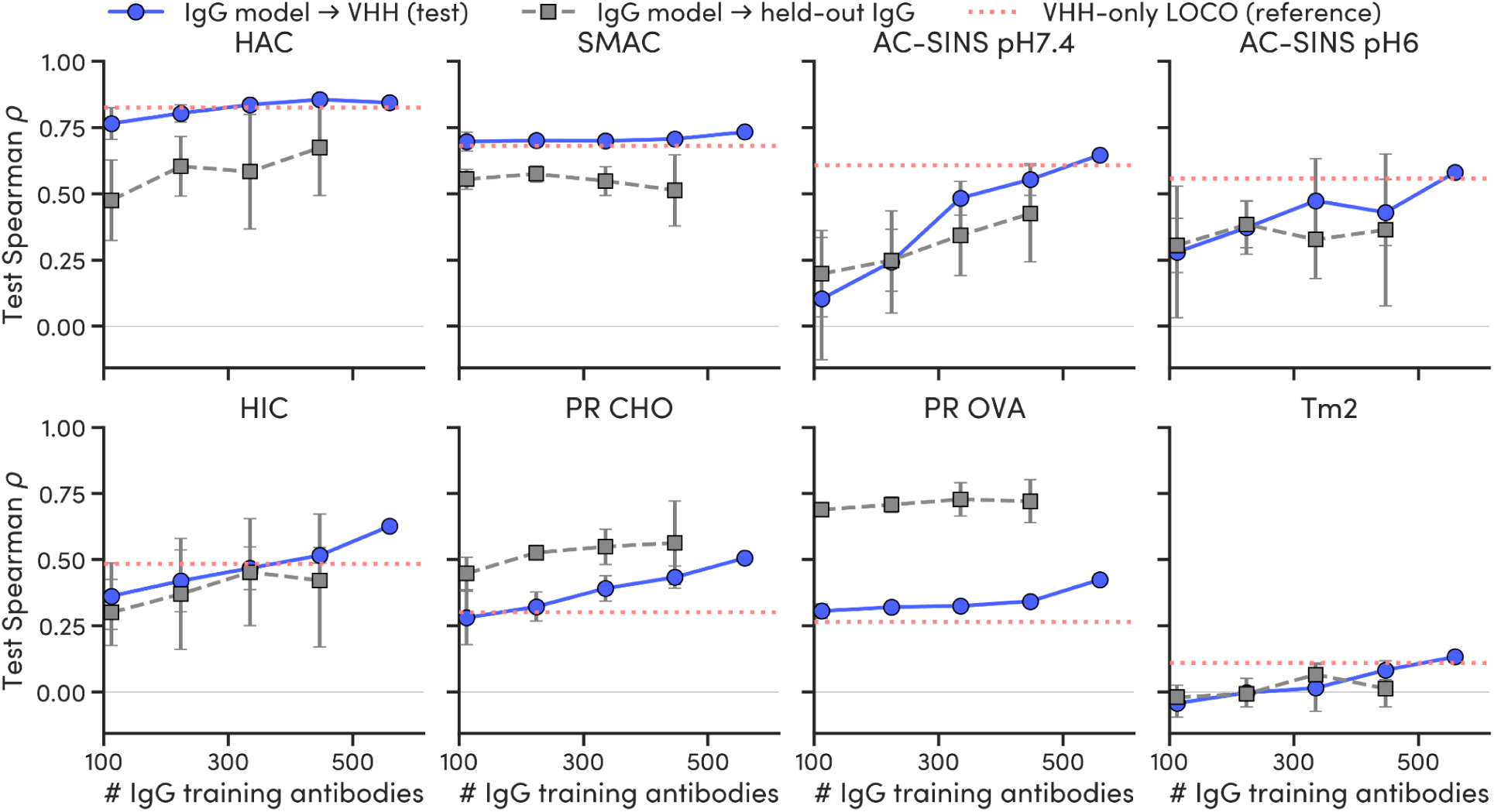
Model performance scaling for cross-format for each property. Both performance on held-out VHH-Fcs and held-out IgGs are shown. Error bars represent the standard deviation of the cross-validation strategy

GDPa5 is released as a public resource alongside GDPa1 [Arsiwala et al., 2025], GDPa3 [van Niekerk et al., 2026], and GDPa4 [Ritter et al., 2026], building a growing suite of standardized, multi-format developability datasets for the community. Together, these datasets, models, and the cross-format framework presented here establish a practical, data-efficient path for developability prediction in emerging antibody formats, one that leverages the IgG data already generated by the field in tandem with new, large-scale format-specific campaigns for every new modality.

Moving forward, the ceiling of predictive fidelity will continue to be dictated by data volume and quality, highlighting the enduring utility of automated, high-throughput platforms. While off-the-shelf tabular neural networks like TabICLv2 and TabPFN performed well, there remains a significant opportunity in aligning and pre-training tabular models explicitly for the task of protein property prediction. Models natively designed to integrate high-dimensional sequence embeddings with biophysical priors and auxiliary assay cascades will likely further transition antibody developability from an empirical screening bottleneck into a fully tractable computational discipline.

## Materials and Methods

### Library Design and Sequence Selection

VHH sequences were curated from PLAbDAb-nano [Gordon et al., 2025], a database of annotated single-domain antibody sequences. The full database was filtered to remove sequences fewer than 100 amino acids in length and to deduplicate identical entries, yielding a pool of 2,306 candidate sequences. From this pool, 160 sequences were selected using farthest-point sampling (FPS) seeded with 36 clinical nanobody sequences drawn from Gordon et al. [Gordon et al., 2026]. FPS iteratively selects sequences that maximize the minimum pairwise distance to all previously selected points in the ESM2 embedding space [Lin et al., 2023], ensuring maximal diversity relative to the clinical seeds while preserving coverage of clinically relevant sequence space. Selected VHH sequences were formatted as VHH-Fc fusions using a human IgG1 Fc region to confer FcRn-mediated half-life extension and enable Protein A purification.

### Antibody Production and Purification

VHH-Fc antibodies were expressed via transient transfection in Chinese Hamster Ovary (CHO) cells and purified by single-step Protein A affinity capture, following protocols established for the PROPHET-Ab platform [Arsiwala et al., 2025]. Post-purification quality metrics including yield, concentration, endotoxin, and purity were recorded for each sample. All 160 antibodies were profiled for the full developability panel; however, the modeling analyses were restricted to VHH-Fcs with SEC monomer purity ≥80% (n = 100).

### Biophysical Assays

GDPa5 was profiled across 10 developability assays on the PROPHET-Ab high-throughput platform: size-exclusion chromatography (SEC, % monomer), hydrophobic interaction chromatography (HIC RT), standup monolayer affinity chromatography (SMAC RT), heparin affinity chromatography (HAC RT), affinity-capture self-interaction nanoparticle spectroscopy (AC-SINS) in PBS pH 7.4 and 20 mM Histidine, 20 mM NaCl, pH 6.0, polyreactivity against CHO cell lysate (PR CHO) and ovalbumin (PR OVA), baculovirus particle (PR BVP) with SuperBlock and BSA blocking buffers, and nanoscale differential scanning fluorimetry (nanoDSF, reporting Tm1 and Tm2). Full assay protocols are described in Arsiwala et al., 2025 and Ritter et al., 2026. All methods were applied without modification.

#### Data Availability

All raw data and sequence information from this study are publicly available and contain:

1. “Definitions” sheet describing each column header corresponding to a data output;
2. “Sequences” sheet documenting the amino acid sequences for all 160 VHH-Fcs produced in this work, the origin of the sequence, and, if it is clinical, the clinical name of the sequence;
3. “Assay Data - average” sheet summarizing average, standard deviations and number of replicates for each developability assessment (one row per antibody).
4. “Assay Data - raw” sheet showing the raw, tidy format developability data;

#### Sequence Featurization

Sequences were featurized using four complementary strategies. Antibody-specific protein language model (pLM) embeddings were generated using AbLang2 [Olsen et al., 2024], a paired antibody language model pre-trained on germline and variable region sequences. General protein pLM embeddings were generated using ESM2 [Lin et al., 2023]. For both pLM models, per-residue embeddings were mean-pooled across sequence positions and reduced to 50 dimensions by principal component analysis prior to model input. Spatial structural features were computed using DeepSP [Kalejaye et al., 2024], which generates surface property descriptors predictive of aggregation and nonspecific binding from sequence. Isoelectric point (pI) was calculated from the VHH sequence using the biopython library. One-hot encodings (OHE) of ANARCI aligned, AHO-numbered [James & Deane, 2016] sequences served as the null hypothesis baseline. For all cross-format experiments, featurizations were applied to IgG heavy chain sequences or VHH sequences only, to match the single-chain nature of the VHH domain.

#### Model Architectures

Four model architectures were evaluated. TabICLv2 [de Freitas, 2025], a tabular in-context learning model, was evaluated in zero-shot (ZS) and fine-tuning (FT) modes. TabPFN v2.5 [Hollmann et al., 2025], a transformer-based tabular foundation model, was similarly evaluated in ZS and FT modes. XGBoost [Chen & Guestrin, 2016] and ridge regression served as gradient-boosted and linear baselines, respectively. For each assay, a full ablation across all architectures and featurization strategies was performed, and the best-performing combination is reported. All models were trained to predict continuous assay values. Performance was evaluated using Spearman’s rank correlation coefficient (ρ).

#### Intra-Format Cross-Validation

Intra-format model performance was evaluated using leave-one-cluster-out cross-validation (LOCO CV). Sequences passing the SEC purity threshold (n = 100) were clustered by sequence identity using mmseqs2 [Steinegger & Söding, 2017], yielding 20 distinct clusters. For the purpose of gathering statistics, singleton clusters were excluded from evaluation. For each LOCO fold, models were trained on all clusters except one and evaluated on the held-out cluster. The pooled Spearman ρ across all held-out sequences is reported as the primary intra-format performance metric.

#### Cross-Format Evaluation

Cross-format models were trained on a 559-antibody IgG dataset: comprising public datasets GDPa1 (n=246) [Arsiwala et al., 2025] and GDPa3 (n=80) [van Niekerk et al., 2026] and additional proprietary Ginkgo internal data (n=288) filtered by the same SEC % monomer threshold as the VHH data. This reduced the amount from 618 to 559. Heavy chain sequences were used as input and full-mAb assay values as prediction targets. PR BVP data were not generated for the IgG training set and are excluded from all cross-format modeling. Trained models were evaluated zero-shot on the GDPa5 VHH-Fc library without any format-specific fine-tuning.

#### Spike-in Analysis

To evaluate whether augmenting IgG training data with VHH-Fc examples improves cross-format performance, increasing proportions of GDPa5 VHH-Fc data (0%, 5%, 10%, 15%, 20%, 25%) were added to the IgG training set. For each spike-in proportion, LOCO CV was applied to the combined dataset. The best-performing model at each proportion was compared to the zero-shot IgG baseline. Full results are shown in **Figure S7.**

#### Auxiliary Feature Cascade

Auxiliary models were constructed by augmenting sequence-only feature vectors with experimental values of selected assay measurements - specifically HIC, HAC, and AC-SINS pH 7.4, identified as the most reliably cross-format predicted properties. For each target assay, all combinations of available auxiliary features were evaluated and the best-performing combination reported. Improvements with Δρ < 0.05 were not considered meaningful, reflecting the practical cost of incorporating additional experimental measurements.

#### Cross-Dataset Comparison

For the 34 VHH sequences present in both GDPa5 and the Gordon et al. dataset [Gordon et al., 2026], rank-order agreement was assessed using Spearman’s ρ. Error bars for GDPa5 measurements represent technical replicates. Error bars for the Gordon et al. dataset are as reported in the original publication.

#### Statistical Analysis and Code Availability

Model performance was quantified using Spearman’s ρ throughout. Data analysis and modeling were carried out in Python 3.12 with scikit-learn, XGBoost, scipy, and standard scientific libraries. The supplementary file describes the modeling summary including the full model ablation study, featurizer ablation, what training dataset and test datasets were used, and the resulting evaluation metrics.

## References

1. Arsiwala A, et al.. A high-throughput platform for biophysical antibody developability assessment to enable AI/ML model training. mAbs. 2025;17(1):2593055. doi:10.1080/19420862.2025.2593055

2. Carter PJ, Lazar GA. Next generation antibody drugs: pursuit of the “high-hanging fruit.” Nat Rev Drug Discov. 2018;17(3):197–223. doi:10.1038/nrd.2017.227

3. Chen T, Guestrin C. XGBoost: a scalable tree boosting system. Proceedings of the 22nd ACM SIGKDD International Conference on Knowledge Discovery and Data Mining. 2016:785–794. doi:10.1145/2939672.2939785

4. De Vlieger, Dorien, et al. “Single-Domain Antibodies and Their Formatting to Combat Viral Infections.” Antibodies, vol. 8, no. 1, 2019, p. 1. 10.3390/antib8010001.

5. Duggan, Sean. “Caplacizumab: First Global Approval.” Drugs, vol. 78, no. 15, 2018, pp. 1639–1642. 10.1007/s40265-018-0989-0.

6. Fernández-Quintero, Monica L., et al. “Assessing Developability Early in the Discovery Process for Novel Biologics.” mAbs, vol. 15, no. 1, 2023. 10.1080/19420862.2023.2171248.

7. Gordon GL, et al. Characterising nanobody developability to improve therapeutic design using the Therapeutic Nanobody Profiler. Commun Biol. 2026. doi:10.1038/s42003-026-09594-y

8. Gordon GL, et al. PLAbDab-nano: a database of camelid and shark nanobodies from patents and literature. Nucleic Acids Res. 2025;53(D1):D535. doi:10.1093/nar/gkae881

9. Grinsztajn, Léo, et al. “TabPFN-2.5: Advancing the State of the Art in Tabular Foundation Models.” arXiv, 2025. https://arxiv.org/abs/2511.08667.

10. Guan, Davy, et al. “Can Tabular In-Context Learners Generalize to Biomolecular Property Prediction?” arXiv, 2026. https://arxiv.org/abs/2606.31126.

11. Hollmann N, et al. Accurate predictions on small data with a tabular foundation model. Nature. 2025. doi:10.1038/s41586-024-08328-6

12. Jain T, et al. Biophysical properties of the clinical-stage antibody landscape. Proc Natl Acad Sci USA. 2017;114(5):944–949. doi:10.1073/pnas.1616408114

13. James L, Deane CM. ANARCI: antigen receptor numbering and receptor classification by sequence. Bioinformatics. 2016;32(2):298–300. doi:10.1093/bioinformatics/btv552

14. Jarasch A, et al. Developability assessment during the selection of novel therapeutic antibodies. J Pharm Sci. 2015;104(6):1885–1898. doi:10.1002/jps.24430

15. Jovèevska I, Muyldermans S. The therapeutic potential of nanobodies. BioDrugs. 2020;34(1):11–26. doi:10.1007/s40259-019-00392-z

16. Kalejaye L, et al. DeepSP: deep learning-based spatial properties to predict monoclonal antibody stability. Comput Struct Biotechnol J. 2024. doi:10.1016/j.csbj.2024.05.029

17. Kraft, Thomas E., et al. “Heparin Chromatography as an In Vitro Predictor for Antibody Clearance Rate Through Pinocytosis.” mAbs, vol. 12, no. 1, 2019. 10.1080/19420862.2019.1683432.

18. Lin Z, et al. Evolutionary-scale prediction of atomic-level protein structure with a language model. Science. 2023;379(6637):1123–1130. doi:10.1126/science.ade2574

19. Mitchell, Laura S., and Lucy J. Colwell. “Comparative Analysis of Nanobody Sequence and Structure Data.” *Proteins: Structure*, Function, and Bioinformatics, vol. 86, no. 7, 2018, pp. 697–706. 10.1002/prot.25497.

20. Muyldermans S. Nanobodies: natural single-domain antibodies. Annu Rev Biochem. 2013;82:775–797. doi:10.1146/annurev-biochem-063011-092449

21. Olsen TH, et al. Addressing the antibody germline bias and its effect on language models for improved antibody design. bioRxiv. 2024. doi:10.1101/2024.02.02.578678

22. Qu J, Holzmüller D, Varoquaux G, Le Morvan M. TabICLv2: a better, faster, scalable, and open tabular foundation model. arXiv. 2026. arXiv:2602.11139

23. Raybould MIJ, et al. Five computational developability guidelines for therapeutic antibody profiling. Proc Natl Acad Sci USA. 2019;116(10):4025–4030. doi:10.1073/pnas.1810576116

24. Ritter S, et al. Decoding bispecific antibody developability: design rules and predictive models from a 160-member library. bioRxiv. 2026. doi:10.64898/2026.06.15.732449

25. Sakhnini, Laila I., et al. “Prediction of Antibody Non-Specificity Using Protein Language Models and Biophysical Parameters.” mAbs, vol. 18, no. 1, 2026. 10.1080/19420862.2026.2678000.

26. Steinegger, M., and J. Söding. “MMseqs2 Enables Sensitive Protein Sequence Searching for the Analysis of Massive Data Sets.” Nature Biotechnology, vol. 35, 2017, 28. pp. 1026–1028. 10.1038/nbt.3988.

27. Svilenov, Hristo L., et al. “Approaches to Expand the Conventional Toolbox for Discovery and Selection of Antibodies with Drug-Like Physicochemical Properties.” mAbs, vol. 15, 2023. 10.1080/19420862.2022.2164459.

28. van Niekerk L, Moller J, Ritter S et al. 2025 Ginkgo Datapoints Antibody Developability Competition Outcomes: Limited Model Performance and A Call for Data Standardization. mAbs. 2026;18(1). doi:10.1080/19420862.2026.2634216

29. Waibl, Franz, et al. “Conformational Ensembles of Antibodies Determine Their Hydrophobicity.” Biophysical Journal, vol. 120, 2021, pp. 143–157. 10.1016/j.bpj.2020.11.010.

30. Xin, Jiayi, et al. “Improved Therapeutic Antibody Reformatting Through Multimodal Machine Learning.” arXiv, 2025. 10.48550/arXiv.2509.19604.

